# The maintenance of centriole appendages and motile cilia basal body anchoring relies on TBCCD1

**DOI:** 10.1101/2023.07.26.549647

**Authors:** Bruno Carmona, Carolina Camelo, Manon Mehraz, Michel Lemullois, Mariana Lince Faria, Étienne Coyaud, H. Susana Marinho, João Gonçalves, Sofia Nolasco, Francisco Pinto, Brian Raught, Anne-Marie Tassin, France Koll, Helena Soares

## Abstract

Centrosomes are organelles consisting of two structurally and functionally distinct centrioles, with the mother centriole having complex distal (DA) and subdistal appendages (SDA). Despite their importance, how appendages are assembled and maintained remains unclear. This study investigated human TBCCD1, a centrosomal protein essential for centrosome positioning, to uncover its localization and role at centrioles. We found that TBCCD1 localizes at both proximal and distal regions of the two centrioles, forming a complex structure spanning from SDA to DA and extending inside and outside the centriole lumen. TBCCD1 depletion caused centrosome mispositioning, which was partially rescued by taxol, and the loss of microtubules (MTs) anchored to centrosomes. TBCCD1 depletion also reduced levels of SDA proteins involved in MT anchoring such as Centriolin/CEP110, Ninein, and CEP170. Additionally, TBCCD1 was essential for the correct positioning of motile cilia basal bodies and associated structures in *Paramecium*. This study reveals that TBCCD1 is an evolutionarily conserved protein essential for centriole and basal body localization and appendage assembly and maintenance. A BioID screening also linked TBCCD1 to ciliopathy-associated protein networks.

## INTRODUCTION

The centrosome is a microtubule (MT) and actin-organizing center in animal cells (Conduit et al., 2015; Farina et al., 2016). It comprises two MT-based centrioles around which the pericentriolar material (PCM), a dynamic matrix of hundreds of proteins, is organized (Woodruff et al. 2014). The PCM provides a structural scaffold to concentrate soluble components of the cytoplasm, like tubulin and γ-tubulin, around the centrioles, thus promoting MT nucleation (Gopalakrishnan et al. 2011, Dictenberg et al. 1998). By nucleating MTs and organizing the cytoskeleton, centrosomes are involved in a multitude of cellular functions, such as the spatial organization of the cytoplasm, cell polarity, cell motility, cell adhesion, and cell division, during which the centrosomes assist in the formation of the mitotic spindle (Lüders and Stearns, 2007). The centrosome is also critical for cilia (Bettencourt-Dias and Glover, 2007), complex MT-based organelles with an axoneme that protrudes from the cell surface. Indeed, the centrioles mature in basal bodies (BBs) to template axonemal growth. Cilia can be either motile or immotile (primary) and perform motility and sensory functions critical for the homeostasis of adult tissues and embryonic development (for review Goetz and Anderson, 2010 and Satir and Christensen, 2007). Cilia absence, abnormal assembly, and malfunction cause human diseases collectively known as ciliopathies (Braun and Hildebrandt, 2017; Mitchison and Valente, 2017).

The centrosome has two structurally and functionally asymmetric centrioles in vertebrate cycling cells. The older (mother) centriole’s distal end is decorated with complex radial structures called distal (DA) and sub-distal appendages (SDA) (Paintrand et al., 1992; Vorobjev and Chentsov, 1982), while the daughter centriole is also enriched in specific proteins, *e.g.,* Centrobin (Gudi et al., 2011; Zou et al., 2005). DA assembly proceeds in early mitosis, starting with the recruitment by CEP83 of its most proximal components, SCLT1 and CEP89. This is followed, in late mitosis, by the SCLT1-mediated recruitment of the most external components FBF1, CEP164, and ANKRD26 (Bowler et al., 2019; Ye et al., 2014; Tanos et al., 2013; Sillibourne et al. 2011). Super-resolution microscopy showed that these proteins are organized in a cone-shaped architecture composed of nine distal appendage blades within an embedded matrix. In the context of cilia, these structures localize to the distal region of the BB around the ciliary gate (Yang et al., 2018).

Studies of DA components’ loss of function support their critical role in both ciliogenesis and ciliogenesis-associated processes (Ye et al., 2014; Joo et al., 2013; Tanos et al., 2013; Reiter et al., 2012; Schmidt et al., 2012; Rosenbaum and Witman 2002). For primary cilia formation, the mother centriole is converted into a BB that migrates to the cell cortex while undergoing a multi-step maturation process where its distal end is remodeled. During this process, DA components mediate the docking of pre-ciliary vesicles, probably derived from the Golgi apparatus, to the mother centriole and then their fusion with the cytoplasmic membrane promoting BB anchoring (Cajanek and Nigg, 2014; Ye et al., 2014; Tanos et al., 2013, Sillibourne et al., 2013; Joo et al., 2013; Schmidt et al., 2012; Graser et al., 2007). Similarly, in multiciliated cells, the DA protein CEP164 is required to recruit, to BBs, proteins Chibby1 (Cby1), FAM92A, and FAM92B, as well as the Rab11-Rab8 axis, which allows the formation of ciliary vesicles and later BB docking (Siller et al., 2017; Lapart et al., 2019). Besides ciliogenesis, some appendages’ components seem to be also involved in cell cycle regulation, DNA damage response, epithelial-to-mesenchymal transition, and apoptosis (Slaats, 2014; Chaki et al., 2012).

Similarly to DA assembly, the assembly of SDA is also hierarchical, and several SDA proteins, including ODF2/Cenexin, CEP128, Centriolin/CEP1/CEP110, CCDC68, CCDC120, Ninein, and CEP170, have been identified and mapped in the centriole structure (Chong et al., 2020; Huang et al., 2017; Mazo et al., 2016; Schrøder et al., 2012; Guarguaglini et al., 2005; Ou et al., 2002; Nakagawa et al., 2001; Mogensen et al., 2000). ODF2 localizes closer to the mother centriole barrel and is a critical factor for the ordered assembly of the other SDA proteins (Chong et al., 2020; Huang et al., 2017; Mazo et al., 2016), with Ninein and CEP170 being the SDA components furthest from the centriole wall and the last to be assembled. SDA assembly is complex and may involve alternative pathways and other components (reviewed in Tischer et al., 2021; Uzbekov and Alieva, 2018).

SDA are important for MT nucleation and anchorage at the centrosome leading to a radial organization of the MT network (Uzbekov and Alieva, 2018; Delgehyr et al., 2005; Guarguaglini et al., 2005; Dammermann and Merdes, 2002; Mogensen et al., 2000; Quintyne et al., 1999). By participating in MT organization, several SDA components (*e.g.,* ODF2 and VFL3/CCDC61) are required in interphase cells for centrosome positioning in close association with the nucleus (Pizon et al., 2020, Hung et al., 2016). Components of SDA have also been shown to localize to the basal foot of motile cilia and to the multiple basal feet of primary cilia, for example, ODF2 (Kunimoto et al.,2012; Ishikawa et al., 2005; Nakagawa et al., 2001), Ninein (Mogensen et al., 2000), CEP170 (Guarguaglini et al., 2005) and Centriolin/CEP110 (Gromley et al., 2003). In multiciliated cells, the basal foot is an electron-dense conical structure that extends from one side of the BB barrel connecting each BB to a subapical MT network (Ohata and Alvarez-Buylla, 2016; Herawati et al., 2016; Kunimoto et al., 2012; Clare et al., 2014). In both cilia types, the basal foot has a modular architecture with three main domains connected by Ninein and Centriolin/CEP110, but the constituent proteins in these domains are differentially organized, probably reflecting cilia functional differences (Nguyen et al., 2020). The basal foot is lost in airway multiciliated cells expressing a mutated *Odf2* gene (Herawati et al., 2016; Kunimoto et al., 2012). This loss is accompanied by a disruption of the polarized organization of apical MT arrays, as well as an incorrect positioning and loss of the polarized alignment of BBs and consequent failure of ciliary beating coordination (Kunimoto et al., 2012). Interestingly, desynchronized cilia beating is also observed when the apical MTs are disturbed by nocodazole (Herawati et al., 2016). The fact that the basal foot and SDA share components but show distinct architectures suggests that SDA proteins may combine/reorganize to assemble the basal foot (Garcia and Reiter, 2016; Uzbekov and Alieva, 2018; Bornens, 2002). The role of the basal foot in organizing MTs is critical for the function of this BB structure and has similarities with the role of SDA in MT anchoring to the centrosome (Kunimoto et al., 2012; Clare et al., 2014).

Our previous work identified a new centrosomal protein, TBCC domain-containing human (TBCCD1), which is related to both TBCC (tubulin cofactor C), a GTPase activating protein for tubulin involved in the tubulin folding pathway, and the GTPase RP2 (retinitis pigmentosa 2) (Gonçalves et al., 2010; Veltel et al., 2008). TBCCD1 depletion in RPE-1 human cells severely affects the position of the centrosome relative to the nucleus, with profound consequences on the integrity of the Golgi, cell size, cell migration, and ability to assemble primary cilia (Gonçalves et al., 2010). TBCCD1 is conserved throughout the eukaryotic lineage and is important for the spatial organization of the cytoplasm. TBCCD1 loss of function in *Chlamydomonas reinhardtii* causes defects in centriole and flagella number, centriole positioning, and spindle orientation (Feldman et al., 2009). In *Trypanosoma brucei*, TBCCD1 depletion results in the disorganization of the bi-lobe structure and a loss of connection between the centriole and the single unit-copy mitochondrial genome (or kinetoplast) (André et al., 2013). More recently, a study in the brown algae *Ectocarpus* showed that TBCCD1 mutants also present increased cell size, modified Golgi architecture, disruption of the MT network, abnormal positioning of the nucleus, and perturbed apical/basal cell identities during development (Godfroy et al., 2017).

In this study, we addressed the functions of TBCCD1 and refined its centriolar localization analysis in human RPE-1 cells. By super-resolution microscopy alone or combined with expansion microscopy, we observed that TBCCD1 localizes in the two centrioles at both proximal and distal regions. At the mother centriole, TBCCD1 shows a complex organization comprised of substructures occurring both inside and outside the centriole lumen and spanning from the DA to the SDA region. Accordingly, TBCCD1 depletion and overexpression assays differentially affect DA protein levels and SDA maintenance/assembly suggesting that a narrow range of TBCCD1 concentrations is critical for the formation and/or maintenance of these structures. Knowing the importance of DA and SDA in ciliogenesis, we further investigated the function of TBCCD1 in multiciliated cells using the ciliated protozoan *Paramecium tetraurelia,* which bears thousands of motile cilia per cell. In agreement with our observations in human mono-ciliated cells, we found that TBCCD1 depletion in *Paramecium* affected BBs’ positioning/anchoring and BBs’ accessory structures assembly, impacting their local orientation relative to the antero-posterior axis of the cell. Finally, we screened TBCCD1 interactors using proximity-dependent biotin identification (BioID) and found that TBCCD1 had a functionally rich interaction environment that supports TBCCD1 roles and places it in the networks of ciliopathy-associated proteins.

## RESULTS

### 1 Centrosome mispositioning caused by TBCCD1 depletion in RPE-1 cells is accompanied by loss of microtubule aster organization, dispersion of centriolar satellites, and altered cell morphology

Previous work from our group identified TBCCD1 as a critical protein for the correct positioning of the centrosome at the cell center in close association with the nucleus (Gonçalves et al., 2010). In interphase cells, centrosome position results from the balance of pushing and pulling forces on the centrosome exerted by MT dynamics, MT-associated proteins including motors (*e.g.,* kinesins and dyneins), as well as specialized cell cortex anchor sites (Odell et al., 2019; Tanimoto et al., 2018; Wu et al., 2011; Kimura and Onami, 2010; Zhu et al., 2010; Dogterom et al., 2005; Burakov et al. 2003). Consequently, the overall MT organization has a pivotal role in the spatial distribution of pushing and pulling forces (Letort et al., 2016; Théry et al., 2006), and changes in this organization will influence the centrosome’s position. Therefore, to go further in the functional characterization of TBCCD1 in centrosome positioning, we investigated the impact of TBCCD1 knockdown on MT arrays’ organization and nucleation. For TBCCD1 depletion in RPE-1 cells, we used the pool of siRNAs previously described by Gonçalves et al. (2010; see also Figure S1). Consistent with our previous results, TBCCD1 depletion dramatically affected centrosome localization, which moved from the center to the cell’s periphery. RPE-1 control cells and TBCCD1-silenced RPE-1 cells were processed for immunofluorescence microscopy (IF) using antibodies against α-tubulin and CEP170 (Figure 1). In Figure 1-I, panels A-D and Aa and Bb, we present IF panels and schematics that illustrate the population of TBCCD1-depleted cells where MTs seemed to retain their association to the displaced centrosomes, as previously noticed (Gonçalves et al., 2010). However, a detailed analysis of stack images at the centrosomal level (Fig 1-I panels a-c) showed that, despite the mispositioned centrosome being able to attach MTs, some unbound MTs, close to the centrosome were clearly visible, suggesting that they had been released (Figure 1-I, panel b-c see arrows). In this case, the radial organization of the aster (symmetric aster) was lost, and MTs frequently tended to accumulate at one side of the nucleus (asymmetric aster) (Figure 1-I, panels A-E, see arrows). In some cells, only a few MTs seemed to remain anchored to the centrosome, with cells narrowing towards one end (Figure 1-I, panel D). This asymmetric accumulation of MTs in TBCCD1-depleted cells had an impact in cell morphology, supporting the notion that MT organization is crucial for cell shape.

**Figure 1.**
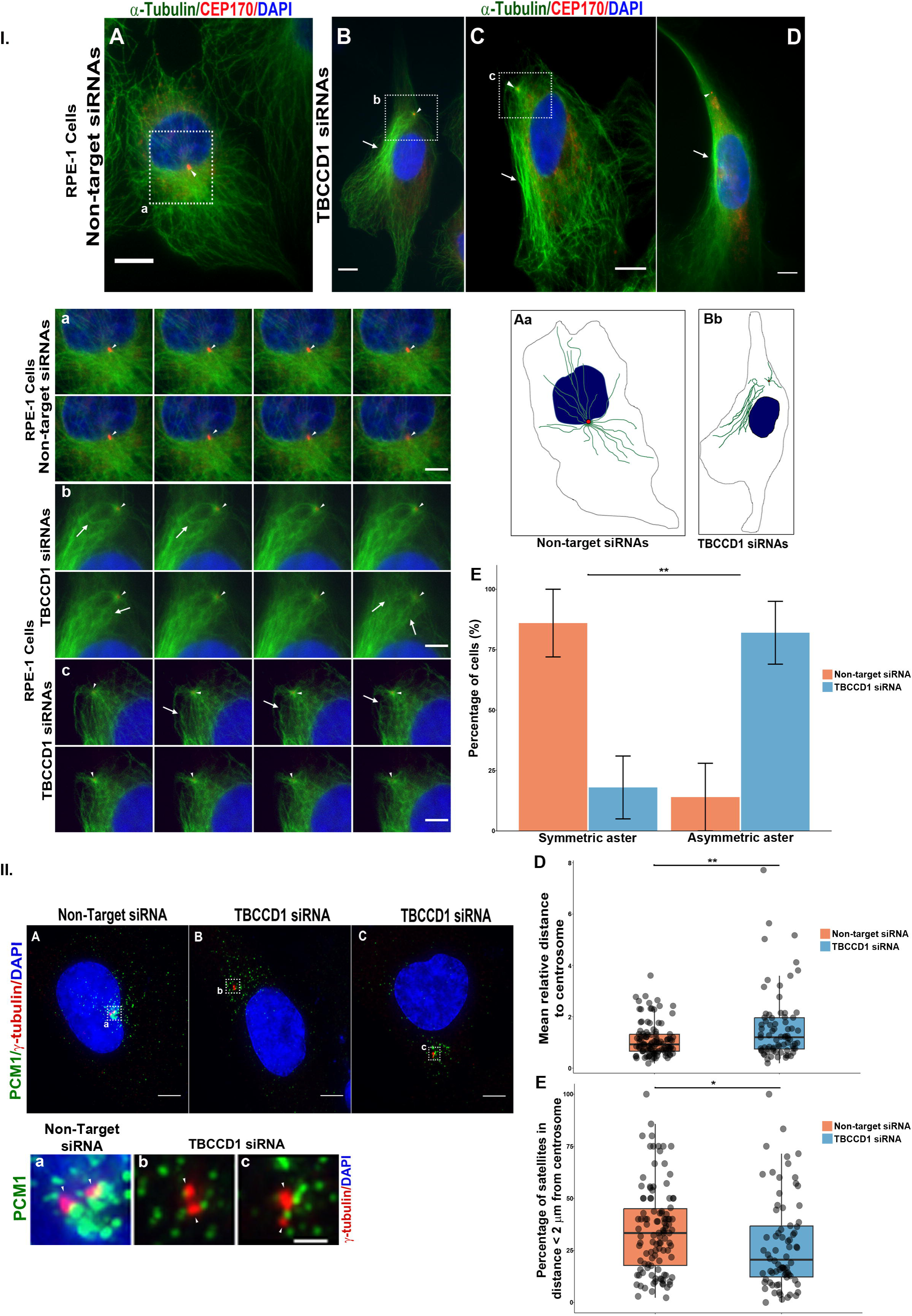
TBCCD1 depletion affects MT cytoskeleton organization and anchoring of MTs to centrosomes causing centriolar satellites dispersion. **I. A-D** Immunofluorescence microscopy images for MT cytoskeleton in RPE-1 cells treated with non-target siRNA and RPE-1 cells where TBCCD1 was depleted by siRNAs (images are representative of three independent experiments). Arrow heads show centrosomes. Arrows show the asymmetric distribution of MTs in TBCCD1 siRNA depleted cells. **Aa and Bb**- Schematic illustration of MT cytoskeleton organization in control and cells treated with TBCCD1 siRNAs. **a-c** - Images from sections of centrosomal details marked by dotted white boxes in A, B, and C, respectively. α-tubulin (green), CEP170 (red), and DNA stained by DAPI (blue). Arrow heads show centrosomes. Arrows show unanchored MTs in the vicinity of the centrosome in TBCCD1 siRNA depleted cells. Scale bar = 10 μm. **E –** Percentage of cells that present a symmetric or asymmetric aster phenotype in RPE-1 cells treated with non-target siRNA cells and RPE-1 cells after TBCCD1 siRNA depletion. Error bars represent 95% confidence interval. Significant differences to the corresponding control are indicated by ** (chi-Square test; p < 0.01). **II. A-C** - Immunofluorescence microscopy images for PCM1 (green) in RPE-1 cells treated with non-target siRNA cells and RPE-1 cells after TBCCD1 depletion by siRNAs. γ-tubulin (red) was used as a centrosome marker, and DAPI was stained in blue for DNA. Scale bar = 5 µm. **a-c** – Centrosome details corresponding to the respective dotted white squares in A-C. Scale bar = 1 µm. **D** – Mean distance of satellites containing PCM1 to the centrosome in RPE-1 cells treated with non-target siRNA and in RPE-1 cells treated with TBCCD1 siRNAs. Two independent experiments were performed with n >100 cells for each condition. Grey dots represent individual measurements, and boxplots are represented in color. Significant differences to the corresponding control are indicated by ** (t-Student test; p < 0.01). **E** – Percentage of centriolar satellites in a distance < 2 µm to the centrosome in RPE-1 cells treated with non-target siRNA and in RPE-1 cells treated with TBCCD1 siRNAs. Two independent experiments were performed with n >100 cells for each condition. Grey dots represent individual measurements, and boxplots are represented in color. Significant differences to the corresponding control are indicated by **** (t-Student test; p < 0.0001).

The quantification of MTs anchored at the centrosome is difficult to evaluate. Moreover, it is well established that centriolar satellites move along MTs toward centrosomes and become dispersed in the cytoplasm upon MT depolymerization (Kubo et al., 1999). Thus, we decided to investigate by IF if depletion of TBCCD1 in RPE-1 cells affected the cellular distribution of centriolar satellites. We hypothesized that if in TBCCD1-depleted RPE-1 cells, centrosomes have less anchored MTs, this should cause a partial dispersion of satellites throughout the cytoplasm, as under MT depolymerization. With this purpose, RPE-1 cells were transfected for 72 h with either a control non-target siRNA or siRNAs specific for TBCCD1. As a marker for the satellites, we used PCM1, a scaffolding protein critical for the assembly and maintenance of centriolar satellites (Prosser and Pelletier, 2020; Hori and Toda, 2017; Dammermann and Merdes, 2002; Kubo et al., 1999) (Figure 1-II). In TBCCD1-silenced cells with misplaced centrosomes, we observed that the satellites were dispersed through the cytoplasm (Figure 1-II, panels A-C and details in a-c). The quantification of the mean distance of the satellites to the centrosome showed that in cells depleted of TBCCD1, the satellites were at a 1.5-fold greater distance from the centrosome than those in control cells (Figure 1-II D). Moreover, if the number of MTs at the centrosome is lower, it would be expected to see less satellites in its neighborhood. In fact, we observed fewer satellites near (< 2μm) the centrosome (34.9% control vs. 27.8% in TBCCD1-RNAi cells) upon TBCCD1 depletion (Figure 1-II E).

Our results strongly support that TBCCD1 loss of function compromised the symmetry of the MT centrosomal aster, causing both a dispersion of satellites in the cytoplasm, and a concomitant decrease in their pericentrosomal localization, supporting the previous evidence that the number of MTs anchored to the centrosome is lower in these cells. Therefore, we decided to investigate if MT stabilization could rescue centrosome positioning in RPE-1 cells silenced for TBCCD1. For this, we used low paclitaxel (taxol) concentrations (0.1 and 0.2 nM) to avoid alterations in cell and nucleus morphology, as well as the formation of MT bundles (Figure S2). RPE-1 cells were transfected with the TBCCD1 siRNA pool for 72 h, and 0.1 nM or 0.2 nM taxol was either added or not (negative control) and 48 h post-transfection for 24 h. Cells were then processed for IF using an antibody against pericentrin to visualize the centrosome, and the centrosome to nucleus distance was assessed as described by Gonçalves et al. (2010). In agreement with our published results (Gonçalves et al., 2010), we found that only 7 % of RPE-1 control cells had the centrosome distanced more than 2 μm from the nucleus, whereas this value increased to 50.7 % in cells depleted of TBCCD1 (Figure 2 III). Noteworthy is the fact that the addition of 0.1 nM taxol partially rescued the phenotype of centrosome displacement to 34.2 %. This rescue was even more pronounced when cells were treated with 0.2 nM taxol, with only 24.5 % of cells showing mispositioned centrosomes. These results demonstrate that the displacement of the centrosome to the periphery of the cell observed in TBCCD1 silenced RPE-1 cells is partially caused by MT destabilization and/or decreased numbers of MT anchored to the centrosome. Interestingly, treatment with taxol also reversed the centrosome mispositioning effect caused by the depletion of the SDA protein ODF2/Cenexin, allowing cells to position their centrosomes at the cell center (Hung et al., 2016). This data indicates that stably anchored MTs at the centrosome are required for MT aster organization and centrosome positioning.

**Figure 2.**
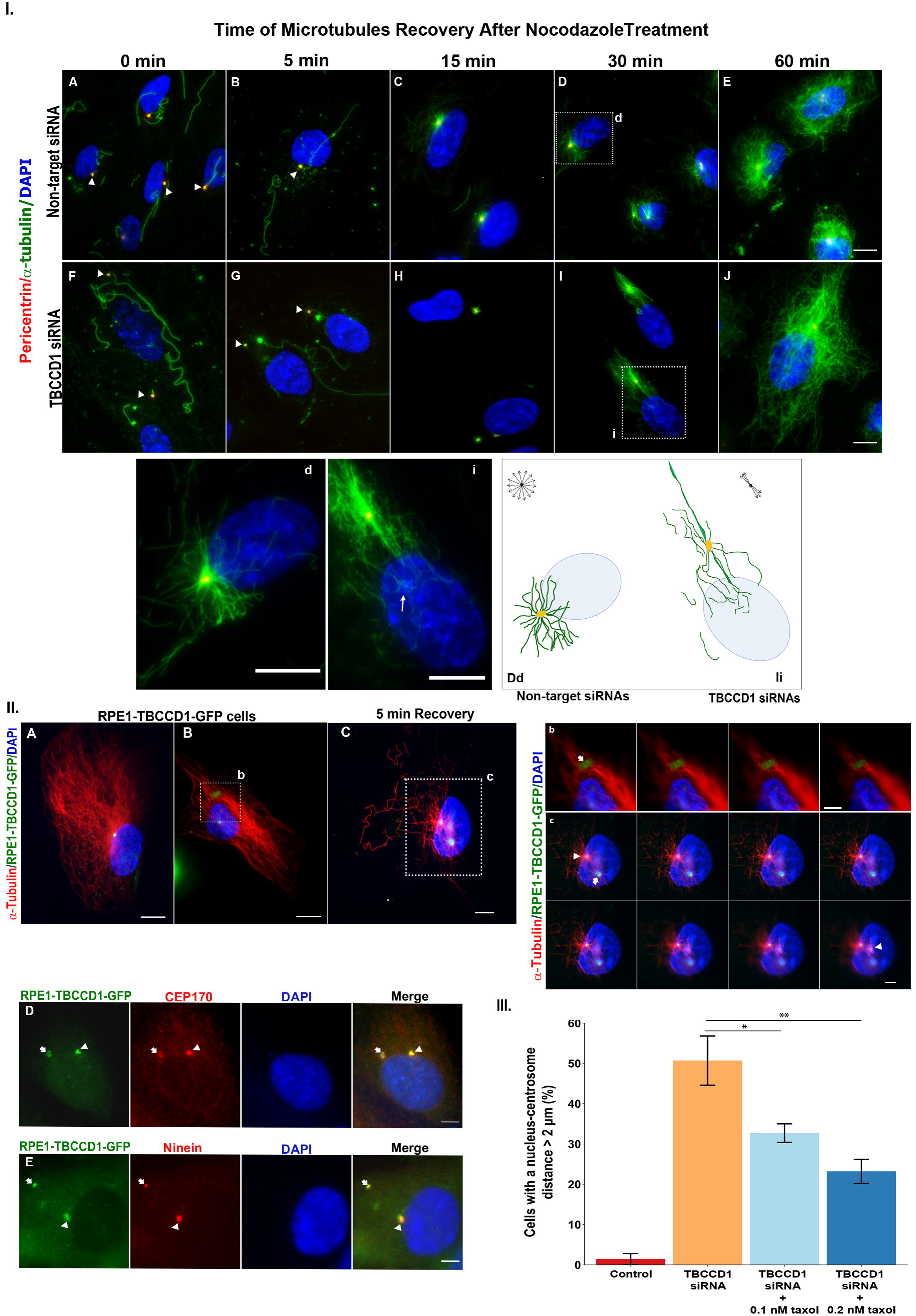
Centrosome mispositioning caused by TBCCD1 siRNAs is partially rescued by paclitaxel (taxol), and TBCCD1 is able to recruit proteins involved in MTs nucleation/anchoring. **I. A-J** - Immunofluorescence microscopy images for tubulin cytoskeleton in RPE-1 cells, in RPE-1 cells treated with non-target siRNA cells and RPE-1 cells depleted of TBCCD1 by siRNAs for 0-, 5-, 15-, 30- and 60-minutes recovery after nocodazole treatment (images are representative of three independent experiments). α-tubulin (green), Pericentrin (red) was used as a centrosome marker, and DNA was stained with DAPI (blue). **d**: detail of recovering MTs in cells treated with non-target siRNA corresponding to the dotted white box area in D. Scale bar = 10 μm. **i** - detail of recovering MTs in TBCCD1 depleted cells showing a preferential distribution along an axis, with MTs growing towards the nucleus, from dotted white box area in I. Scale bar = 10 μm. **Dd-Ii-** Schematic illustration of MT cytoskeleton organization in control and TBCCD1 siRNA depleted cells after recover from nocodazole treatment. **II. A-C** - Immunofluorescence microscopy images for MT cytoskeleton in RPE-1 cells constitutively expressing TBCCD1-GFP. In C cells were recovering for 5 minutes after nocodazole treatment (images are representative of three independent experiments). α-tubulin (red), TBCCD1-GFP (green), and DNA stained with DAPI (blue). Dotted white squares in B and C: detail of recovering MT in RPE-1 constitutively expressing TBCCD1-GFP. **b-c** - Images from sections corresponding to details of recovering MTs in RPE-1 expressing TBCCD1-GFP (B and C). TBCCD1-GFP aggregates (arrows) nucleate MTs after recovery from nocodazole treatment. Arrowheads indicate centrioles. Scale bar = 10 μm except for b and c where scale bar = 5 μm. **D-E -** Immunofluorescence microscopy images for TBCCD1 small aggregates in RPE-1 cells constitutively expressing TBCCD1-GFP stained for centrosomal proteins CEP170 (D) and Ninein (E) (both in red) and DNA is stained with DAPI (blue). Cep170 and Ninein co-localize with TBCCD1 at aggregates, as indicated by arrows. Arrowheads indicate centrioles. Scale bar = 10 μm. **III.** Centrosome mispositioning caused by low levels of TBCCD1 was partially rescued by taxol in a dose-dependent way. Percentage of cells with a centrosome-nucleus distance > 2 μm in control RPE-1 cells and cells treated with siRNAs specific to TBCCD1 in the absence and presence of taxol (0.1 nM and 0.2 nM). Three independent experiments were performed with n > 200 cells. Error bars represent standard deviation. Significant differences to the corresponding control are indicated by * (t-Student test; p < 0.05) or ** (t-Student test; p < 0.01).

The ability of the centrosomes of TBCCD1-silenced RPE-1 cells to nucleate MTs after depolymerization with 30 μM nocodazole (30 min at 4° C) was also tested for different recovery times (0, 5, 15, 30, and 60 min after nocodazole washout). We used antibodies against α-tubulin and anti-pericentrin to visualize MTs and the centrosome (Figure 2-I panels A-J). As previously shown by us, mispositioned centrosomes in TBCCD1-depleted cells were still able to nucleate MTs, and a small aster was already visible 15 min after the nocodazole washout (Figure 2-I, panel H). However, washing out the unpolymerized tubulin decreased the background significantly, allowing us to observe that, contrary to control cells, in which MTs regrew radially, in TBCCD1-depleted cells, MTs recovered and grew in preferential directions (Figure 2-I, panel I/i and scheme Ii). Growing MTs from displaced centrosomes tended to orient along the axis formed by the centrosome and the nucleus. Interestingly, some of these MTs were in close contact with the nucleus (Figure 2-I, panel i). We may wonder if this subset of MTs also exists in control cells. An attractive hypothesis would be that a subset of MTs could provide information to the centrosome regarding its position/distance to the nucleus through the forces generated via MTs anchored to the nuclear envelope. Misplaced centrosomes positioned away from the nucleus helped to unmask this subset of the MT population (Figure 2-I, panels A-J and schemes Dd and Ii). These experiments suggested that TBCCD1 may be involved in MT anchoring and organization at the centrosome. In this context, we decided to explore the RPE-1 cell line constitutively overexpressing TBCCD1-GFP (see Figure S1 for cell line characterization) (Cardoso et al., 2014), since that cell line frequently displays TBCCD1 aggregates of different sizes, probably due to moderate overexpression levels of TBCCD1-GFP. A striking feature of these TBCCD1 aggregates was that after nocodazole treatment followed by its washout, they were able to nucleate and grow MTs (see Figure 2-II, panels C/c). In panel c of Figure 2-II, we show that these aggregates were in a distinct plane to that of the two centrioles that are separated, probably due to nocodazole treatment, and MTs growing from the aggregate were radially organized. Moreover, some acentrosomal aggregates were also detected in cells not subjected to nocodazole treatment. Similarly, MTs seemed to be captured/anchored by/to these acentrosomal aggregates (Figure 2-II, panel B/b). Remarkably, TBCCD1 aggregates were also stained by antibodies against CEP170 and Ninein (Figure 2-II panels D and E), two proteins that localize to SDA and the proximal end of both centrioles (Mazo et al., 2016). We do not have evidence that TBCCD1 is able to bind to MTs. However, at the end of cell division, TBCCD1-GFP was recruited from the cytoplasm to the midzone, decorating small segments of spindle MTs in this region (Figure S1 IIC; III and Smovie1). Together, these results suggest that TBCCD1 can recruit proteins involved in MTs nucleation/anchoring.

### 2 TBCCD1 localizes at the proximal and distal ends of centrioles in RPE-1 cells

Our results showing that: (i) MTs stabilization by taxol can partially rescue the mispositioning of centrosomes in TBCCD1-depleted cells; (ii) TBCCD1 loss affects MT aster organization and anchoring to centrosomes; (iii) acentrosomal aggregates containing TBCCD1 observed in overexpressing cells can nucleate/anchoring MTs; prompted us to do a detailed localization analysis of TBCCD1 in the centrosome. For this, we used super-resolution microscopy 3D-Structured Illumination Microscopy (SIM) and several centriole markers: (i) the DA protein CEP164 (Graser et al., 2007), and (ii) the SDA proteins ODF2, CEP128, Centriolin/CEP110 (Mazo et al., 2016; Gromley et al. 2003), CEP170 (mature mother centriole marker in G1 cells; Guarguaglini et al., 2005) and Ninein (Delgehyr et al., 2005; Mogensen et al., 2000) (Figure 3). To identify TBCCD1, we used a polyclonal serum against TBCCD1 in RPE-1 cells, the RPE-1 cell line constitutively expressing TBCCD1-GFP, and in both cases anti-α-acetylated tubulin as a marker of centriolar MTs (Figure 3). After MT depolymerization with cold shock (4° C), the anti-TBCCD1 sera showed that TBCCD1 localized, independently of MTs, at both ends of one of the centrioles, whereas in the other centriole, it localized mainly at one end (Figure 3 A and B). In the centriole with TBCCD1 at both ends, the protein originated a partial discrete ring above the centriolar cylinder at one of the ends connected by small arms to a bright dot inside the lumen of the centriole (Figure 3 Aa and Bb, see arrowheads). At the opposite end of this centriole, we can also observe the presence of TBCCD1 (Figure 3A/a and B/b, concave arrowheads). In the other centriole, a simpler structure containing TBCCD1 was also observable at one end (Figure 3A/a and B/b, arrow).

**Figure 3.**
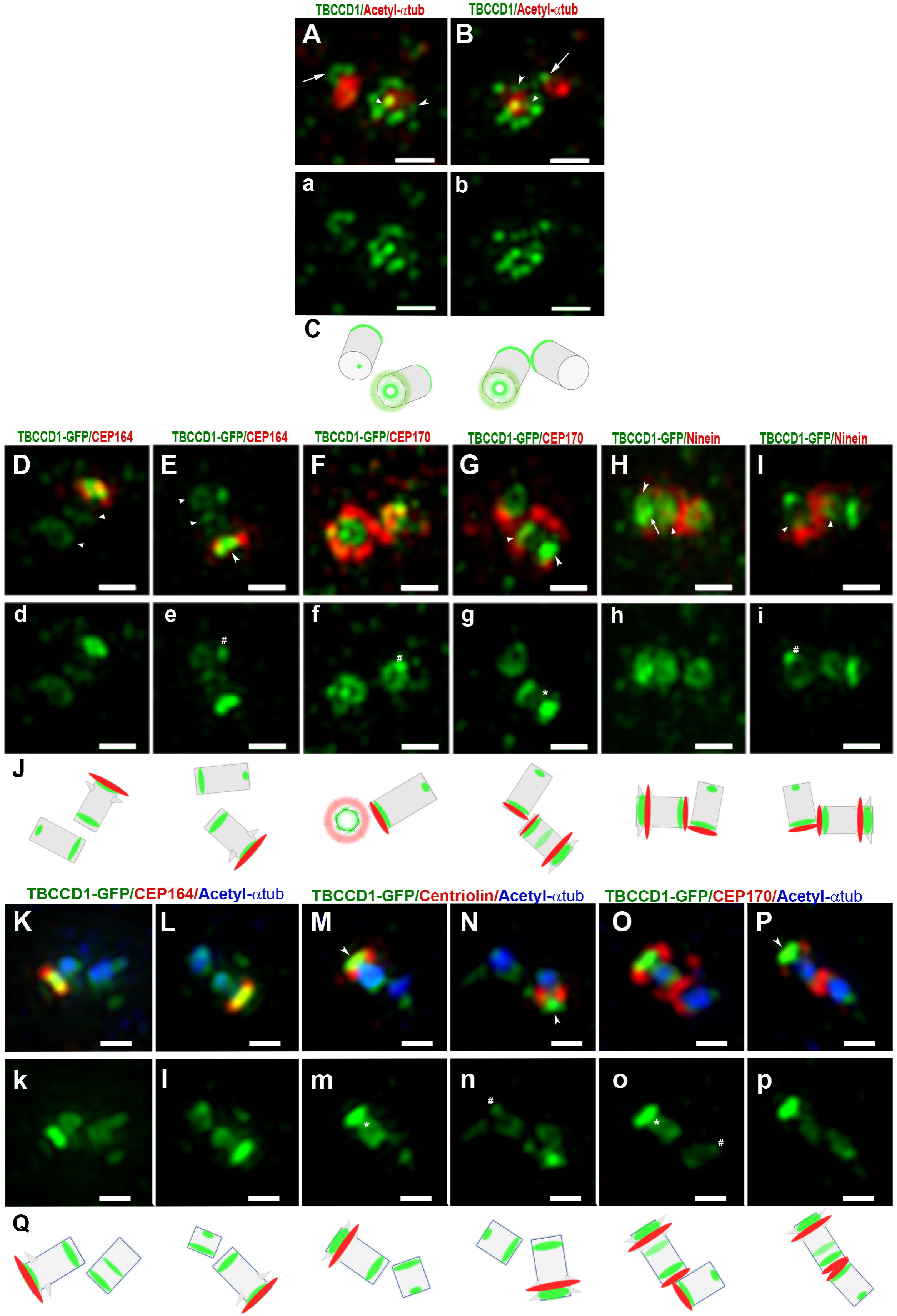
Localization of TBCCD1 in centrosomes of RPE-1 cells. **Aa-Bb-** Z projections of super-resolution microscopy (3D-SIM) images of RPE-1 cells, where MTs were depolymerized with cold shock (4° C), stained with an antibody directed against TBCCD1 (green), an antibody against acetylated-α-tubulin (red) to show centriolar MTs. TBCCD1 localized at both ends of one of the centrioles, whereas in the other was localized mainly at only one end (see arrows). In one of the centrioles ends TBCCD1 originated a partial discrete ring above the centriole walls with a strong dot at the center. The two structures seemed to be connected by small arms (see arrowheads). Concave arrowheads indicate TBCCD1 localization in the opposite centriole end. **C**- Schematic representations of the TBCCD1 (green) localization and the centrioles’ orientation. **Dd-Ii and Kk-Pp**- RPE-1 cells constitutively expressing TBCCD1-GFP (green) were stained with antibodies directed against the DA protein CEP164 (red; D, E, K, and L), SDA proteins CEP170 (red; F, G, O and P), Ninein (red; H, I) and Centriolin/CEP110 (red; M, N). An antibody against acetylated-α-tubulin was used to show centriolar MTs (blue, K-P). SDA proteins CEP170 and Ninein are also localized at centrioles’ proximal ends. In RPE-1 cells constitutively expressing TBCCD1-GFP, the localization pattern of TBCCD1 was similar to that revealed by the antibody specific to TBCCD1. The centriolar markers show that at the mother centriole’s most distal end, TBCCD1 localization corresponded to a complex structure. In E, G, H, M, N, and P, this structure shows a longitudinal section resembling a cone where the vertex points to the proximal end of the mother centriole (concave arrowheads). This structure seemed to be composed, in the most distal region, by a ring (F and H) at the level of the DA protein CEP164 (D, E, K, and L) connected to a smaller ring/dot corresponding to the cone’s vertex (H, arrow) at the level of the SDA proteins Centriolin/CEP110 (M and N), CEP170 (G, O, and P) and Ninein (H and I). In a few mother centrioles, a third very faint TBCCD1 open ring could be detected at the centriole midzone (G, M, and O; asterisks). TBCCD1 was also organized into one ring that localized near the mother centrioles’ proximal end. In the daughter centriole, TBCCD1 originated a ring at the centriole proximal end (Dd-Ii and Kk-Pp), and an intense TBCCD1 dot was frequently observed in the opposite end (E, I, N, and O; see hashes). Scale bars = 0.5 μm. **J and Q**- Schematic representations of the TBCCD1-GFP (green) localization and the centrioles orientation. Other centriole markers are in red. Results are from 3 independent experiments.

In RPE-1 cells constitutively expressing TBCCD1-GFP, the localization pattern of TBCCD1 was similar to that revealed by the anti-TBCCD1 sera in RPE-1 cells. However, the fluorescence intensity was higher in the different localizations (Figure 3 D-I and K-P). We performed a localization analysis with multiple markers: CEP164 (DA), Centriolin/CEP110 (SDA), Ninein, and CEP170 (SDA and proximal end). The use of these markers allowed us to conclude that the structure described in Figure 3A and B as a ring connected to a dot inside the centriole lumen localizes at the distal end of the mother centriole (Figure 3 D/d-I/i). In fact, at this localization, TBCCD1 was present in a complex structure (Figure 3 panels D-I and K-P) that spanned from the inside to the outside of the centriole lumen and showed a longitudinal section resembling a cone with the vertex pointing to the proximal end (Figure 3, panels E, G, H, M, N and P; see concave arrowheads). At its most distal region, this structure is composed by a ring connected to a smaller ring/dot probably corresponding to the cone’s vertex (Figure 3 H, see arrow). The most distal ring (base) of the cone localized at the level of the DA protein CEP164 (Figure 3 D/d, E/e, K/k, and L/l), whereas the vertex localized at the level of the SDA proteins Centriolin/CEP110, Ninein, and CEP170 (Figure 3 G/g, H/h, M/m, N/n, O/o, P/p). In this analysis, we consistently observed that TBCCD1 was also organized into one ring localized near the proximal end of the mother centriole (Figure 3 panels D/d-I/i and K/k-P/p). In a few mother centrioles, a third very faint TBCCD1 open ring could be detected at the centriole midzone (Figure 3, panel G/g, M/m, and O/o; see asterisk). Regarding the daughter centriole, TBCCD1 formed a ring at the proximal end, and a bright TBCCD1 dot was frequently observed in the opposite end (see Figure 3, E, F, I, N, O; see hash).

Due to the complexity of TBCCD1’s pattern of localization and to add additional detail to it within the centriolar structure, we combined super-resolution SIM with expansion microscopy (Gambarotto et al., 2019). This approach increases the resolution by a factor of ∼ 4 compared to conventional super-resolution microscopy. It confirmed the observations made with super-resolution microscopy alone and added additional details on the structures where TBCCD1 localizes in both centrioles. Unfortunately, the anti-TBCCD1 sera did not recognize the TBCCD1 protein after the procedures of the expansion protocol. Consequently, all the results were obtained using the constitutively expressing TBCCD1-GFP cell line. Super-resolution expansion microscopy shows two distinct structures at the most distal region of the mother centriole. However, it revealed specific details of these structures that were not visible before. The TBCCD1 ring at the most distal end of the centriole described above (Figure 3) was revealed to be a punctuated ring with a diameter of 315 ± 47 nm (38 analyzed centrosomes), localizing to a region consistent with that of DA (Figure 4 A/a-C/c, and E; details in stacks A1/a1-A2/a2 and C1/c1-C2/c2; see arrows). This ring was composed of discrete structures linked by arms to a conical inner structure (see Figure 4 A and in the group of stacks of A and C (Figure 4 A1/a1-A2/a2; C1/c1-C2/c2). The cone wall was also composed of individual structures, like umbrella rods, linking the base of the cone to the vertex, a small ring/spot (see Figure 4 panel A1/a1 and panel C1/c1 and illustration in A1’). Inside the centriole lumen, this conical structure had a base diameter of 83 ± 13 nm and a height of 80 ± 8 nm (Figure 4A and C and scheme E). Details of this complex structure can be observed in stack images of Figure 4 A1-A2 and 4 C1 and C2 and are illustrated in A1’. For a few mother centrioles (7.9 % of 38 analyzed centrosomes), we observed TBCCD1 embedded in the centriole MT wall that frequently had small projections toward the outside of the centriole barrel (see Figure 4 D, see asterisks). This localizes at 104 ± 15 nm of the distal end of the centriole acetylated MT wall, which is the expected region for SDA localization (Figure 4 D; see asterisks). A TBCCD1 ring of 186 ± 22 nm was observed at the proximal end of the mother centrioles (Figure 4 A, B, and D, see also stacks A1/a1-A2/a2). In the case of the daughter centriole, the pattern of TBCCD1 localization was less complex since TBCCD1 mainly localized at the proximal end as a ring of 171 ± 23 nm (see Figure 4 A, B, and D) and at the distal end as a variable structure of 90 ± 36 nm (for example Figure 4 A and D, concave arrowheads).

**Figure 4.**
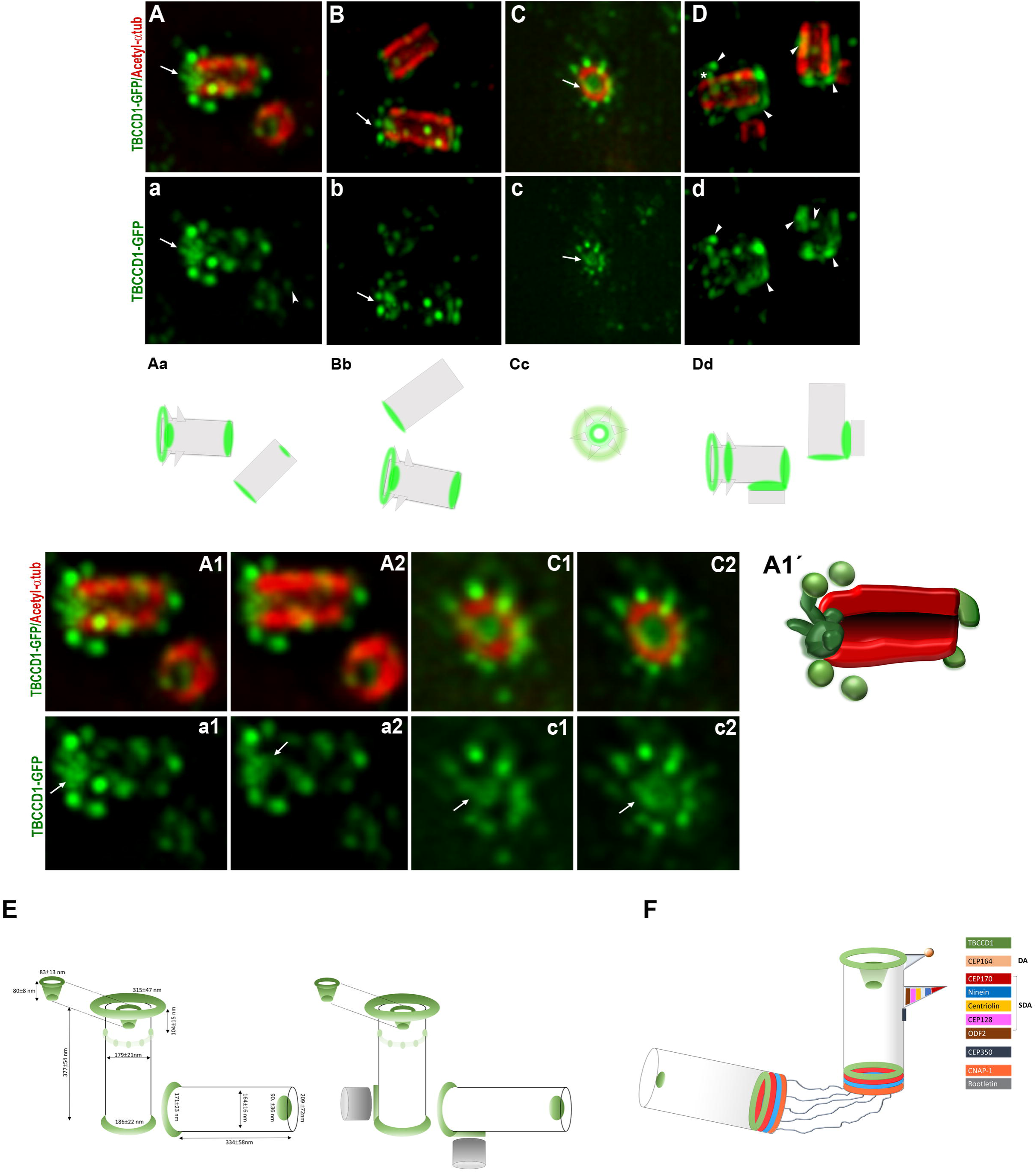
Super-resolution SIM expansion microscopy localization of TBCCD1. **A-D-** RPE-1 cells constitutively expressing TBCCD1-GFP (green) were expanded using a super-resolution SIM expansion combined technique and stained for acetylated-α-tubulin (red), showing the centriole wall. Super-resolution expansion microscopy confirms the presence of TBCCD1 in two distinct structures at the most distal region of the mother centriole. Maximum intensity Z projections show the organization of TBCCD1: TBCCD1 at the distal region of the mother centriole was a punctuated ring that localized at a region consistent with DA (A-D); This ring was composed of discrete structures linked by some arms to a conical inner structure (A-C; A1/a1-A2/a2 and C1/c1-C2/c2). The conical structure comprised individual structures, like umbrella rods, that converge from the base of the cone to the vertex, a small ring/spot (A, B, and C; see arrows). Details of this complex structure can be observed in section images of A1/a1-A2/a2 and C1/c1-C2/c2. A schematic representation of this structure is in A1**’**, where TBCCD1-GFP is in green, and the centriole wall is in red. Centrosomes without any of the TBCCD1 distal and proximal structures were never observed. In the daughter centriole, the pattern of TBCCD1 localization was less complex since TBCCD1 mainly localized at the proximal end as a ring (for example, in A and B) and at the distal end as a variable dot structure (A and D; concave arrowheads). In duplicating centrosomes, TBCCD1 mainly maintained its pattern of localization in both centrioles (D/d) but also localized outside the centriolar cylinder wall in the region where procentrioles were assembled and close to their proximal end (D/d). Scale bars = 0.5 μm. **E** - Schematic representation summary of major centriolar localization of TBCCD1-GFP and respective sizes corresponding to average ± standard deviation of 38 analyzed centrosomes from 3 independent experiments. **F -** Schematic representation of TBCCD1 localization relative to several DA, SDA, and centriole proximal region markers (see also Figure 3).

In duplicating centrosomes, TBCCD1 mainly maintained its localization pattern in both centrioles. However, it concentrated in the outside of the centriolar cylinder wall, in the region where procentrioles were being assembled and close to their proximal end (Figure 4 D/d arrowheads), suggesting that TBCCD1 was required for procentriole assembly.

Together, our results showed that TBCCD1 was localized in a complex structure at the distal region of the mother centriole, composed of a ring at the level of the DA protein CEP64 connected to a conic structure in the lumen of the centriole, extending from the centriole tip to the SDA’ region (Centriolin/CEP110, CEP170, Ninein). Moreover, similarly to Ninein and CEP170, TBCCD1 was found at the proximal region of both centrioles, and our observations show that during the cell cycle, TBCCD1 accumulates at the proximal region of procentrioles possibly playing a role in their assembly.

### 3 Tightly regulated levels of TBCCD1 are required to maintain the centriole distal region

Considering TBCCD1’s localization pattern at both centrioles and the fact that TBCCD1 depletion affects MT anchoring to the centrosome and MT network organization, we proceeded to examine the functional relationship of TBCCD1 and DA and SDA. For this, TBCCD1 was depleted in RPE-1 cells by RNAi, and markers for distinct centriolar domains were examined using wide-field microscopy and 3D structured illumination microscopy (SIM) alone or combined with expansion microscopy whenever better resolution was required to clarify the results. TBCCD1 knockdown did not affect the fluorescence intensity, organization, or localization of the proximal centriolar end protein C-NAP1 at the centrosome (Figure 5-I, panels L, M, and N), as well as of the DA protein CEP164 (Figure 5-I, panels E-G, and N; Figure S3). This indicates that TBCCD1 is dispensable for the recruitment of these proteins to the centriole proximal end and to DA, respectively. Also, TBCCD1 depletion did not affect the organization and localization of the SDA components, ODF2 and CEP128 (Figure S3 panels C, D, and E, F, respectively), localizing closer to the centriole wall (Mazo et al., 2016).

**Figure 5.**
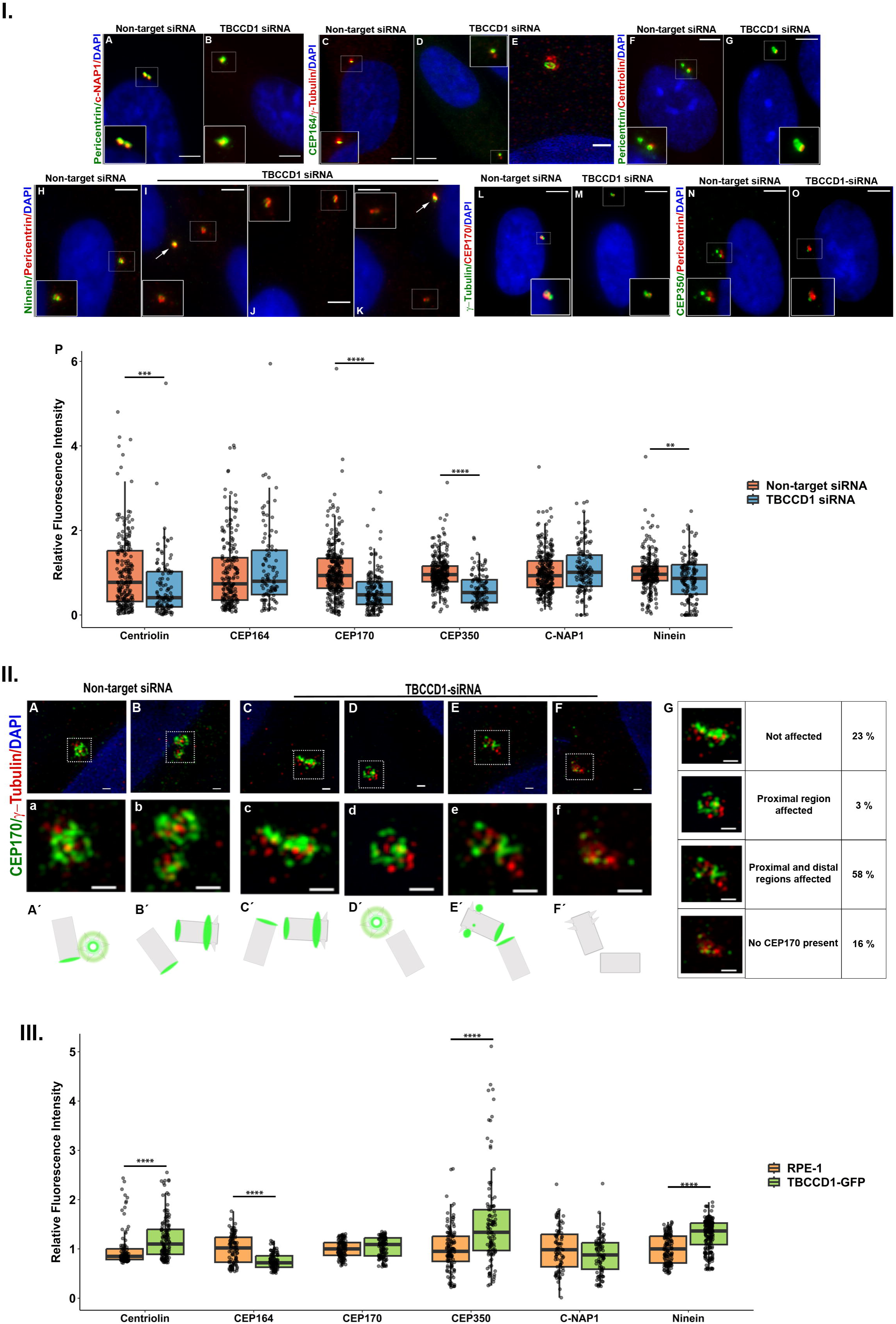
TBCCD1 depletion compromises the centriolar levels of Centriolin/CEP110, Ninein, and CEP170 involved in centrosome MT-anchoring and also of CEP350, whereas TBCCD1 overexpression affects the levels of CEP164 as well asCentriolin/CEP110, Ninein, and CEP350. **I -** Immunofluorescence microscopy images of RPE-1 cells treated with non-target siRNAs and TBCCD1 specific siRNAs. **A-B** – Impact of TBCCD1 depletion in C-NAP1 (red). Pericentrin (green) was used as a centrosome marker, and DAPI stained DNA in blue. **C-E** – Impact of TBCCD1 depletion in CEP164 (green). γ-tubulin (red) is used as a centrosome marker, and DAPI stains DNA in blue. **E** – image obtained using Super-Resolution 3D SIM with a scale bar = 1 μm. **F-G** – Impact of TBCCD1 depletion in Centriolin/CEP110 (red). Pericentrin (green) was used as a centrosome marker, and DAPI stained DNA in blue. **H-K** –Impact of TBCCD1 depletion in Ninein (green). Pericentrin (red) was used as a centrosome marker, and DAPI stains DNA in blue. **L-M** – Impact of TBCCD1 depletion in CEP170 (red). γ-tubulin (green) was used as a centrosome marker, and DAPI stains DNA in blue. **N-O** – Impact of TBCCD1 depletion in CEP350 (green). Pericentrin (red) was used as a centrosome marker, and DAPI stained DNA in blue. In all images, inserts represent zoom images of centrosomes surrounded by white dotted squares. As expected, TBCCD1 depletion causes the mispositioning of centrosomes that are displaced towards the periphery of the cell (Gonçalves et al., 2010). Arrows indicate centrosomes with a nucleus-centrosome distance similar to control cells. Scale bars = 5 μm. **P** - Relative fluorescence intensity for centriolar proteins (Centriolin/CEP110, CEP164, CEP170, CEP350, C-NAP1, and Ninein) in TBCCD1 siRNA experiment in RPE-1 in comparison to RPE-1 treated with non-target siRNA. Fluorescence intensity was measured using Cell Profiler v4.2.1. Three independent experiments were performed. Grey dots represent individual measurements. Boxplot represented in colors. Significant differences to the corresponding control are indicated by ** (t-Student test; p < 0.01), *** (t-Student test; p < 0.001), and **** (t-Student test; p < 0.0001). **II.** Super-Resolution 3D SIM images for CEP170 in RPE-1 cells subjected to non-target siRNAs and TBCCD1 specific siRNAs. **A-F** -Impact of TBCCD1 depletion in CEP170 (green). γ-tubulin (red) was used as a centrosome marker, and DAPI stains DNA in blue. **a-f** – Zoom images of the respective centrosome images. Scale bars = 500 nm. **Á-F’**- Schematic illustration of centrioles orientation and CEP170 staining. **G –** Percentage of the different phenotype classes observed for CEP170 protein in TBCCD1 depleted cells. **III.** Relative fluorescence intensity of DA protein CEP164; SDA proteins Centriolin/CEP110, CEP170 (also in centrioles proximal end) and Ninein (also in centrioles proximal end); centriolar protein CEP350 and centrioles proximal end protein C-NAP1 in RPE-1 cells constitutively overexpressing TBCCD1-GFP in comparison to RPE-1 cells. Fluorescence intensity was measured using Cell Profiler v4.2.1. Three independent experiments were performed. Grey dots represent individual measurements. Boxplot represented in colors. Significant differences to the corresponding control are indicated by **** (t-Student test; p < 0.0001).

In the case of Centriolin/CEP110, we observed a 34 % decrease in its fluorescence intensity at the centrosome (n =113 centrosomes) in the TBCCD1 depletion background (Figure 5-I J, K, and N). This was accompanied by a decrease of 14 % in the fluorescence intensity of Ninein (n=148 centrosomes), which was almost absent in 9.5 % of the centrosomes (Figure 5-I, panels A-D and N). In cells with low levels of TBCCD1, the most affected component of SDA was the protein CEP170 (Figure 5-II, panels E-H). The fluorescence intensity of CEP170 at the centrosome decreased by 43 % (n=159 centrosomes) compared to the centrosomes of control cells (Figure 5-I N). A more detailed analysis of TBCCD1 loss of function using 3D SIM revealed four main phenotypic classes based on how severely CEP170 localization was affected (n = 31 centrosomes), namely: CEP170 not/slightly affected (23%) (Figure 5-II E); CEP170 affected only at the proximal end (3%) (Figure 5-III F); CEP170 affected at the distal and proximal localization in both centrioles (58%) (Figure 5-II G); CEP170 staining completely lost (16%) (Figure 5-II, H). For example, panel H of Figure 5-II displays a cell where the distal ring organization in the mother centriole was entirely lost, and the remaining CEP170 protein was apparently dissociated from the centriole. The results show that TBCCD1 was critical to maintain CEP170 at the SDA and the proximal ends of centrioles. Finally, we also studied the impact of TBCCD1 RNAi in the fluorescence intensity and localization of the centriolar protein CEP350, which is required for the stability and formation of DA and SDA (Karasu et al., 2022), as well as for MT anchoring (Yan et al., 2006). Our results showed that TBCCD1 depletion caused a 39% decrease in CEP350 levels at the centrosome (n=119 centrosomes).

Prompted by these results, we also investigated the impact of TBCCD1 overexpression on DA (CEP164), on SDA (Centriolin/CEP110, Ninein, and CEP170), and the other centriolar proteins previously analyzed (C-NAP1 and CEP350). For this, we compared the IF staining intensity of the referred centrosomal markers in parental RPE-1 cells with that obtained in the RPE-1 cell line constitutively expressing TBCCD1-GFP (Figure 5-III). In the context of TBCCD1 overexpression, the IF signals at individual centrosomes of Centriolin/CEP110, Ninein, and CEP350 proteins were higher than in WT cells by 21% (n=137 centrosomes), 29% (n=218 centrosomes) and 54 % (n=111 centrosomes), respectively (Figure 5-III). Noteworthy, a 25 % decrease in the levels of CEP164 (n=103 centrosomes) was observed, whereas the levels of CEP170 and C-NAP1 did not change significantly (Figure 5-III).

Our data support a model where the downregulation and/or overexpression of TBCCD1 differentially affects components of SDA and the centriolar protein CEP350. Interestingly, the levels of the SDA protein CEP170 were not affected by higher concentrations of TBCCD1 which were able to affect the DA CEP164 levels. Moreover, tightly regulated levels of TBCCD1 were required for the correct localization/recruitment of the outer group of SDA proteins Centriolin/CEP110, Ninein, and CEP170, but seemed not to be required for those SDA proteins localized closer to the centriole wall, such as ODF2 and CEP128. Therefore, TBCCD1 is required for the stability of the SDA’s more external module, which is involved in MT-anchoring to the centrosome. This is supported by the fact that TBCCD1 depletion and overexpression differentially affected the abundance of CEP350 at the centrosome. CEP350 depletion causes loss of MT anchoring and acute MT network disorganization (Yan et al., 2006), which may explain why displaced centrosomes due to TBCCD1 depletion lose MTs, disperse satellites, and presented affected aster organization as observed in Figure 1.

### 4 TBCCD1 is required for BB positioning/anchoring, assembly and orientation of BB accessory components, and proper cilia beating in Paramecium

In multiciliated cells, newly assembled centrioles are transported toward the cell apical surface, where they reorganize, and as they migrate, they are converted into BBs by the addition of rootlet and basal foot appendages (Boutin et al., 2019). MTs connected to the basal foot of each BB (that localizes opposite the striated rootlet) link the individual BBs to their neighbors and promote the correct positioning and local coordination of cilia orientation (Herawati et al., 2016; Vladar et al., 2012; Werner et al., 2011). In a previous study, we showed that TBCCD1 localizes at the basal body of primary cilia and the basal bodies of motile cilia (Gonçalves et al., 2010). We also showed that TBCCD1-depleted cells are less efficient in primary cilia assembly (Gonçalves et al., 2010), however, the role of TBCCD1 in multiciliated cells was still unclear. Since TBCCD1 is required for the correct assembly of SDA, and the basal foot assembly requires SDA proteins like ODF2 (Kunimoto et al. 2012), we investigated the function of TBCCD1 in a multiciliated cell model. For this, we chose *Paramecium tetraurelia*. This highly differentiated unicellular organism has an elaborated cortical pattern where thousands of BBs (about 4,000 in a typical *Paramecium*, Iftode, and Fleury-Aubusson, 2003) are regularly arranged in longitudinal rows that extend from the anterior to the posterior pole of the cell. Most of the *Paramecium* BBs assemble motile cilia that beat coordinately. Since *Paramecium* BBs, despite some differences, are both structurally and molecularly conserved with the BBs of other eukaryotes, this organism is an excellent model for studying motile cilia.

Two genes encoding TBCCD1 were identified in the macronuclear genome of *Paramecium* (PTET.51.1.P0100138 and PTET.51.1.P0590071 references in *Paramecium* DB https://paramecium.i2bc.paris-saclay.fr/). The amino acid sequences of the proteins encoded by these genes share an identity of 92.1% and a similarity of 98.4%. Both proteins display the expected functional domains of TBCCD1 (Figure S4), the TBCC domain that is localized between the amino acid residues 235 to 349, and the overlapped CARP domain (between the amino acid residues 232 to 303)- found in the conserved family of Cyclase-associated proteins (CAP), actin-binding proteins that regulate actin dynamics (Ono, 2013) (Figure S4). The amino acid sequence of both *Paramecium* TBCCD1 proteins is 24.04 % identical to that of human TBCCD1.

To investigate the localization of TBCCD1 in *Paramecium*, we expressed a full-length TBCCD1-GFP protein in wild-type cells. For this, *Paramecium* cells were transformed by injection in the macronucleus of the DNA construct that expresses TBCCD1 tagged with GFP at the C-terminal end under the control of the calmodulin promoter. TBCCD1-GFP localized at BBs stained with an antibody against poly-glutamylated tubulin (ID5) (Figure 6-I, panel A-C). This localization was also observed in cortical units containing one or two BBs, a distinctive feature of the *Paramecium* cortex (Figure 6-I, panels A and C). In addition, this approach led to the accumulation of the tagged protein at the feeding vacuoles (Figure 6-I, panel A). To improve TBCCD1-GFP localization in *Paramecium,* avoiding the excess background caused by the feeding vacuoles, the cortex of *Paramecium* cells expressing TBCCD1-GFP were purified and processed for confocal microscopy analysis (Figure 6-I, panel D-F). In addition, improving the quality of the TBCCD1-GFP localization, this approach confirmed that TBCCD1 was undoubtedly localized at the *Paramecium* BBs when they were singlets or doublets (see Figure 6-I, panels Fa, and Fb). This observation also reinforced the idea that the association of TBCCD1 with BBs is conserved in the multiple diverse phylogenetic species studied so far, from unicellular to multicellular organisms (André et al., 2013; Gonçalves et al., 2010; Feldman et al., 2009), except for the brown algae *Ectocarpus* (Godfroy et al., 2017).

**Figure 6.**
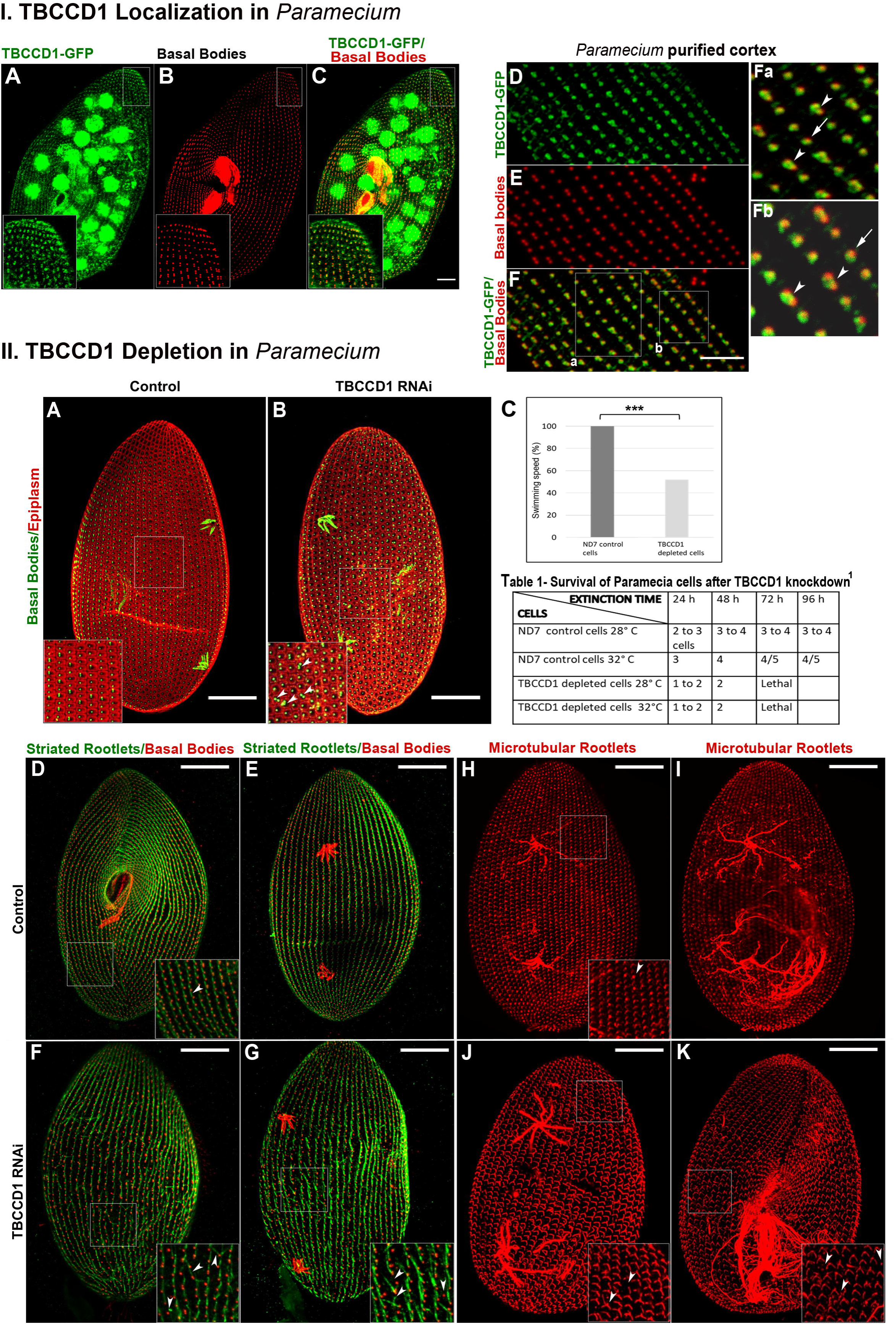
In *Paramecium* cells, TBCCD1-GFP localizes at BBs of motile cilia, and its depletion causes mislocalization of BBs and abnormal striated and microtubular rootlets. **I-** TBCCD1-GFP localization. **A-C -** Z projections of confocal immunofluorescence sections corresponding to the ventral surface of an interphase *Paramecium* transformant expressing TBCCD1-GFP. *Left insets,* the TBCCD1-GFP (green) was localized at BBs stained with an antibody against poly-glutamylated tubulin (red; ID5). The TBCCD1-GFP accumulates at the feeding vacuoles. Scale bars = 10 μm**. D-F-** To reduce the staining background, the cortex of *Paramecium* cells expressing TBCCD1-GFP was purified and processed for confocal microscopy analysis. Dotted white boxes in F correspond to Fa and Fb details showing that TBCCD1 is undoubtedly localized at the *Paramecium* BBs when they are singlets (arrows) or doublets (concave arrowheads). Scale bars = 10 μm. **II-**TBCCD1 depletion phenotypes -BBs localization and epiplasm organization. **A and B** – Z projections of confocal immunofluorescence sections corresponding to the dorsal surface of the cortex of an interphase *Paramecium* (A) and a *Paramecium* cell depleted of TBCCD1 (B), both stained with an antibody against poly-glutamylated tubulin to show BBs (green; ID5) and an antibody against epiplasmins (red; CTS32) that decorates the epiplasm. Dotted white boxes in A and B correspond to left insets showing that in TBCCD1-knockdown cells, the epiplasm units tended to lose their regular hexagonal shape, being rounded or squarer, and many of them contained abnormal numbers of BBs, namely groups of three and four BBs and some BBs are mispositioned/oriented accordingly to the anterior-posterior axis of the cell (concave arrowheads). These phenotypes were never observed in wild-type cells (control). **C** - TBCCD1-depleted cells, 48 h after RNAi treatment, showed a 48% decrease in swimming speed, suggesting defects at the BBs/cilia. Two independent experiments were performed with 80 cells analyzed for each condition. Significant differences to the corresponding control are indicated by *** (t-Student test; p < 0.001). BBs accessory structures- **D-G-** Z projections of confocal immunofluorescence sections corresponding to the ventral (D) and dorsal (E) surface of the cortex of interphase *Paramecium* and dorsal surface of the cortex of *Paramecium* cells depleted of TBCCD1 (F and G), all stained with an antibody specific to striated/kinetodesmal fibers (green; KD4) and with the ID5 antibody to stain BBs (red). Dotted white boxes in D and F-G correspond to the right insets. In cells depleted of TBCCD1, striated/kinetodesmal fibers were misaligned along the antero-posterior axis, pointing to different directions, causing the disruption of the rows of BBs (concave arrowheads). The BBs were also mispositioned. **H-K-** Z projections of confocal immunofluorescence sections corresponding to the dorsal (H, I) surface of the cortex of interphase *Paramecium* cells, and dorsal (J) and ventral (K) surface of the cortex of *Paramecium* cells depleted of TBCCD1, all stained with an antibody against acetylated tubulin (red; TEU318) revealing the BB accessory structures microtubular rootlets. Dotted white boxes in H and J-K correspond to the right insets. The TBCCD1-depleted cells showed BBs with just one microtubular rootlet or without any of these structures (concave arrowheads). Scale bars = 20 μm. ^1^In Table 1, the surviving evaluation was performed by analyzing over time (24, 48, 72, and 96 h after TBCCD1 RNAi) drops initially containing one cell.

Therefore, we next examined the impact of the knockdown of both TBCCD1 genes in *Paramecium*, using the RNAi feeding method according to Galvani and Sperling (2002). To silence simultaneously both TBCCD1 genes, *Paramecium* cells were fed with a mix of bacteria producing dsRNA (see Figure S4), covering 80.5% (1188 nt/1476 nt) of each gene. The efficiency of the dsRNAs to specifically inactivate the targets was evaluated by analyzing the fluorescence intensity of TBCCD1-GFP expressing cells after TBCCD1 depletion in comparison with control cells (Figure S4 III). In RNAi-treated TBCCD1-GFP expressing cells, the GFP signal was lost, and the protein was absent from BBs stained with the antibody against poly-glutamylated tubulin (ID5) (Figure S4 III). These results indicate an almost complete depletion of TBCCD1-GFP. *Paramecium* TBCCD1-depleted cells displayed a 48% decrease in swimming speed 48 h after RNAi treatment, indicating abnormalities at the basal bodies/cilia (Figure 6-II, panel C, Bouhouche et al., 2022). A decrease in cell division rate and death at 72 h are also observed, indicating that TBCCD1 is essential for cell survival (Table 1). Based on these results and the localization of TBCCD1, we investigated the impact on the structure of BB-containing cortical units. For this, cells fed with TBCCD1 dsRNAs were stained for poly-glutamylated tubulin and also with an antibody against epiplasmins (the monoclonal antibody CTS32) that decorates the epiplasm, a submembrane skeleton that is a thin dense layer segmented into regular units where BBs are implanted, originating a hexagonal pattern (Aubusson Fleury et al., 2013; Lynn, 2008) (Figure 6-II, panel A). As previously described (Tassin et al., 2016, Bouhouche et al., 2022), *Paramecium* cells are characterized by different fields displaying BB units containing either one or two BBs or a mix where both types of units coexisted. Remarkably, in TBCCD1-knockdown cells, the epiplasm units tended to lose their regular hexagonal shape, being rounded or squarer (Figure 6-II, panel B). Moreover, 1.9 % ± 1.1 % (n= 10 cells analyzed) of them contained abnormal numbers of BBs, namely groups of three and four BBs, something never observed in wild-type interphase cells (Figure 6-II, panels A and B). In a few units containing BB doublets, the BBs no longer showed the longitudinal alignment along the anterior-posterior axis found in control cells but adopted a more perpendicular orientation to the axis, suggesting defective BB positioning (see Figure 6II, panel B arrowheads) as observed in FOPNL/For20, centrin 2, centrin3, Cep90, and Ofd1 knock down paramecium cells (Aubusson-Fleury et al. 2017; Ruiz et al. 2005; Bengueddach et al., 2017; LeBorgne et al., 2022) Also, an excess of cortical units c.ontaining BB pairs [20.3 % ± 6.7 % in depleted cells (n = 10) vs. 11.6 % ± 2 % in control cells] was also observed upon TBCCD1 depletion as in VFL3-A depletion (Bengueddach et al., 2017). The *Paramecium* BBs contain three major appendages: one long anterior striated rootlet (striated fiber or kinetodesmal fiber) and two microtubular ribbons/rootlets (see scheme in Figure 7 K) (Iftode and Fleury-Aubusson, 2003). These structures help to segregate, guide, and position nascent BBs, maintaining the position of the old ones and, finally, anchor and coordinate them at the cell cortex (Tassin et al., 2016; Jerka-Dziadosz et al., 2013). BB-associated rootlets, both in *Tetrahymena* and *Paramecium*, have been shown to segregate, guide and position the nascent BB (Nabi et al., 2019; Bengueddach et al., 2017; Jerka-Dziadosz et al., 2013). Therefore, we investigated if TBCCD1 silencing impacted the organization of striated rootlets, using the antibody against striated/kinetodesmal fibers (KD4) (Figure 6-II D-G). In control cells, BB-associated structures have a specific orientation relative to the cell’s antero-posterior axis, creating a BB asymmetry and a local intrinsic polarity that is required for correct synchronization of cilia beating (for review Soares et al., 2019, Bouhouche et al., 2022). This organization creates the idea of continuous longitudinal fibers, where BBs are regularly spaced in the ciliate cortex (Figure 6-II D-G). In TBCCD1-depleted cells, the striated rootlets were misaligned along the antero-posterior axis, pointing to distinct directions and originating a complete disruption of the rows of BBs in certain places (Figure 6-II F and G). A closer analysis of the cell cortex showed that some BBs either did not have a striated rootlet or just had a tiny one (4.2 % ± 1.1 %) (Fig 6II F and G; see concave arrowheads). In other cases, some BBs displayed more than one striated rootlet (5.2 % ± 1.2 %) (Fig 6-II F and G; concave arrowheads). The observed defects in striated rootlets were also accompanied by defects in the other BB accessory structures, *i.e.,* the microtubular rootlets (observed in 29 % ± 0.9 %) (Fig. 6-II panels H-K). Using an antibody against acetylated tubulin (TEU318), we observed, either on the dorsal or ventral sides of the cells, BBs with just one microtubular rootlet (11.7 % ± 1.7%), without any of these structures (8.2 % ± 2 %) or with 3 or more (9.1 % ± 0.6 %) (Figure 6-II J-K). These abnormalities involving BBs’ appendages was never observed in control cells, suggesting that they could be the origin of the loss of the BB-specific anterio-posterior orientation and their defective positioning.

**Table 1.**
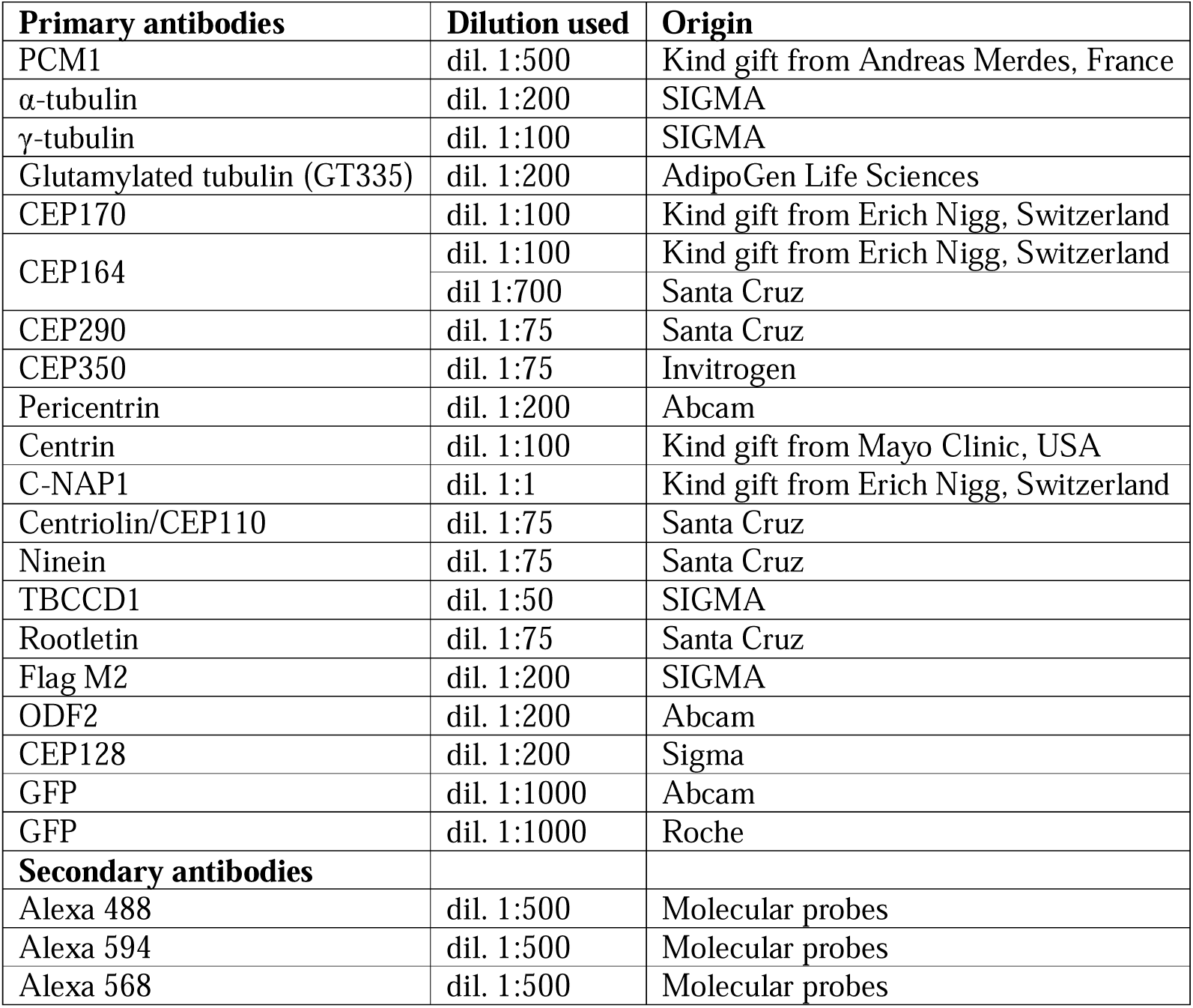
Antibodies and dilutions used human cells.

**Figure 7.**
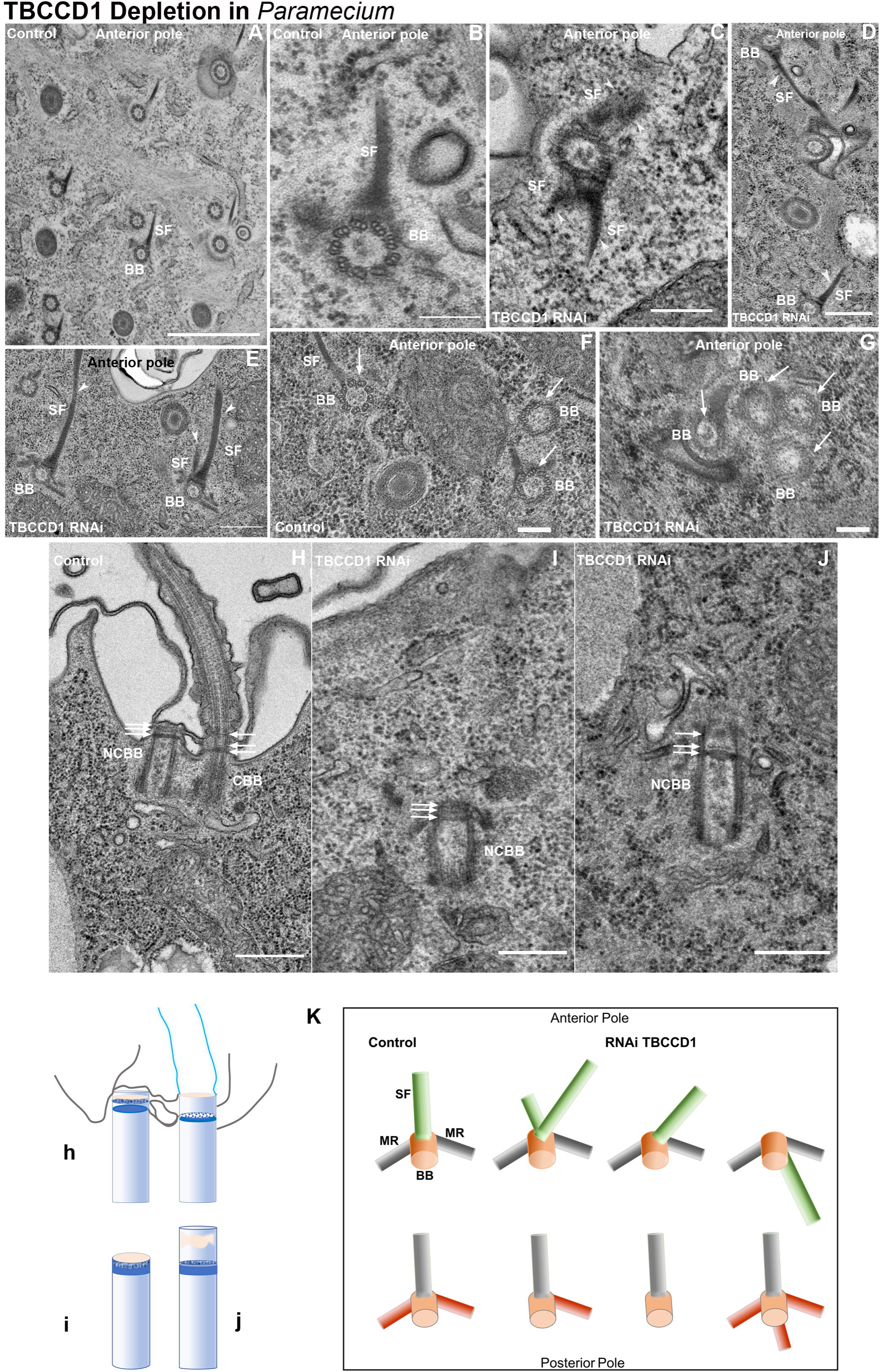
TBCCD1 depletion affects BB number, docking, and accessory structures. **A-B-** Electron microscopy cross-section of wild-type cell: only one striated/kinetodesmal fiber (SF) is associated with BB, all pointing in the same direction to the anterior pole of the cell (A and B). **C-E** - In TBCCD1 depleted cells by RNAi electron microscopy cross sections show extra numbers of striated fibers per BB with altered length (C and E), and these structures are also misaligned in relation to the anterior-posterior axis of the cell (C and D, see arrowheads). **F** - In wild-type cells, cortical units contain one or two BBs aligned with the anterior-posterior axis of the cell. Arrows indicate BBs (BB). **G -** A cortical unit with an abnormal number of BBs (four) lost the antero-posterior orientation, presenting a randomized position. Arrows indicate BBs (BB). **H-J -** In *Paramecium*, anchored BBs are not always ciliated. The BBs that harbor a mature axoneme display a fully organized transition zone composed of three plates: the terminal plate, the intermediate plate, and the axosomal plate occurring adjacent to the ciliary necklace. Electron microscopy longitudinal sections of anchored non-ciliated basal bodies (NCBB) and ciliated basal bodies (CBB) in wild-type (H) and non-ciliated basal bodies (NCBB) in TBCCD1 depleted cells (I, J). The three plates present in the pro-transition zone and the transition zone (H) are indicated by arrows. Undocked (I and J) BBs in TBCCD1 depleted cells a distal end structure resembling that of the non-ciliated anchored BBs (I), whereas others had a distal structure resembling that of the anchored ciliated BBs of wild-type cells (J). Therefore, these BBs appear fully assembled though they are not anchored. **h-j.** Schematic illustration of the observed alterations in distal ends of unanchored BBs of cells depleted of TBCCD1 (i, j) compared to those of wild-type cells (h). **K**- Schematic illustration summary of the defects in *Paramecium* BBs associated structures observed in TBCCD1 depleted cells compared to those observed in wild-type cells. In **A** scale bar = 1 μm; in **B** scale bar = 0.2 μm; in **C** scale bar = 0.25 μm**, D** scale bar = 0.5 μm; **E** scale bar = 0.5 μm, **F** scale bar = 0.2 μm, **G** scale bar = 0.2 μm, in **H-J** scale bar = 0.5 μm.

Next, to strengthen these observations, we conducted an ultrastructural analysis by electron microscopy of BBs/cilia at the cell cortex in cells depleted of TBCCD1 (Figure 7). This analysis confirmed our previous IF observations (Figure 6-II see panels A and B). In cross-sections, cortical units with abnormal BB numbers were identified. For example, the group of four BBs in panel G lost their antero-posterior orientation, presenting a randomized position (Figure 7, compare panel F with panel G).

Regarding the BB accessory structures, the depletion of TBCCD1 led to extra numbers of striated rootlets per BB with altered length. In panel E, we can see two striated rootlets of different lengths emanating from a single BB, whereas a single striated rootlet was always observed in wild-type control cells (compare Figure 7 A and B with Figure 7 E, see scheme K). An abnormal number of short striated rootlets can also be observed in Figure 7 C. Misaligned, very long striated rootlets with wrong antero-posterior orientation were also observed (Figure 7 compare panel A with D; see scheme K).

In *Paramecium*, anchored BBs are not always ciliated, which allows the comparison of the structure of both non-ciliated and ciliated BBs in the same cell (Tassin et al., 2016). Thus, BBs that harbor a mature axoneme display a fully organized transition zone composed of three plates: the terminal plate, the intermediate plate, and the axosomal plate occurring adjacent to the ciliary necklace (see Figure 7, panel H, CBB). The terminal plate extends to link to the epiplasm (Tassin et al., 2016). The non-ciliated anchored BBs (NCBB) were already composed of three plates, although the transition zone is less expanded (see Figure 7, panel and scheme H/h) compared to the transition zone of an anchored ciliated BB. One cell division upon TBCCD1 knockdown, we observed newly assembled BBs that were unanchored (Figure 7, panels and schemes I/i and J/j). Nevertheless, these non-anchored BBs had a transition zone resembling the non-ciliated anchored BBs present in the wild-type cells (Figure 7-panel J/j), appearing to be correctly assembled. This observation indicates that either they were already anchored and that TBCCD1 depletion caused their release from the membrane, or their transition zone was fully maturated in the cytoplasm.

The *Paramecium* experiments were critical to elucidate that TBCCD1 was required for correct positioning, orientation, and anchoring of motile cilia BBs, which parallels its involvement in centrosome positioning and primary cilia assembly in RPE-1 cells. These depletion phenotypes result in an aberrant number of BBs, length and antero-posterior orientation of striated fibers, and numbers and orientation of microtubular rootlets, resulting in a BB mispositioning. These observations resemble the phenotypes observed when components of SDA are depleted from vertebrate multiciliated cells, which are also required for the organization of primary and motile BBs’ accessory components as the basal foot (Kunimoto et al., 2012; Clare et al., 2014). Our results support the notion of a functional relationship between SDA and BBs’ accessory structures and establish TBCCD1 as a new critical protein for the maintenance and assembly of these structures.

### 5 TBCCD1 localizes at the distal end of the BB and recruits Rootletin to the centrosome of the human primary cilium in RPE-1 cells

The localization of TBCCD1 at the proximal and distal regions of the mother centriole of human cells and the results in *Paramecium* showing that TBCCD1 was required for BB positioning, orientation, and anchoring, led us to investigate in more detail TBCCD1 localization at the BBs of primary cilia in RPE1 cells. For this, we assessed if TBCCD1 was maintained at the BB distal region near the transition zone of primary cilia. As a transition zone marker, we used an antibody against CEP290, a protein required for the assembly of this region, where it tethers the axonemal MTs to the ciliary membrane (Wu et al., 2020; Chih et al., 2012; Garcia-Gonzalo et al., 2011; Sang et al., 2011). Ciliogenesis was induced in RPE-1 cells constitutively expressing TBCCD1-GFP by serum-starvation (see Material and Methods), and the cilia axoneme and BB were stained with an antibody against polyglutamylated tubulin except in experiments to assess cilia length, where an antibody against acetylated α-tubulin was used. We observed that TBCCD1-GFP localizes between the distal end of the BB and the base of the axoneme of primary cilia (Figure 8-I panels A-O and scheme P), where it partially co-localized with CEP290 without affecting its levels (Figure 8-III panels E-I). Interestingly, TBCCD1-GFP overexpressing cells showed a decrease of 77 % of their capability to assemble cilia compared to RPE-1 parental cells (Figure 8-II, panel D). Moreover, cilia assembled in cells overexpressing TBCCD1 have an average length of ∼50 % longer (4.75 µm) than that observed in wild-type cells (3.22 µm) (Figure 8-II, panels A-C, E). We also investigated the impact of TBCCD1 siRNAs in the levels of CEP290 in RPE-1 cycling cells and observed that TBCCD1 depletion causes a 60% decrease in CEP290 levels (n=106) (Figure 8-IV E-G). Therefore, these observations identified TBCCD1 at the distal end of the BB extending to the cilia transition zone and affecting cilia length regulation.

**Figure 8.**
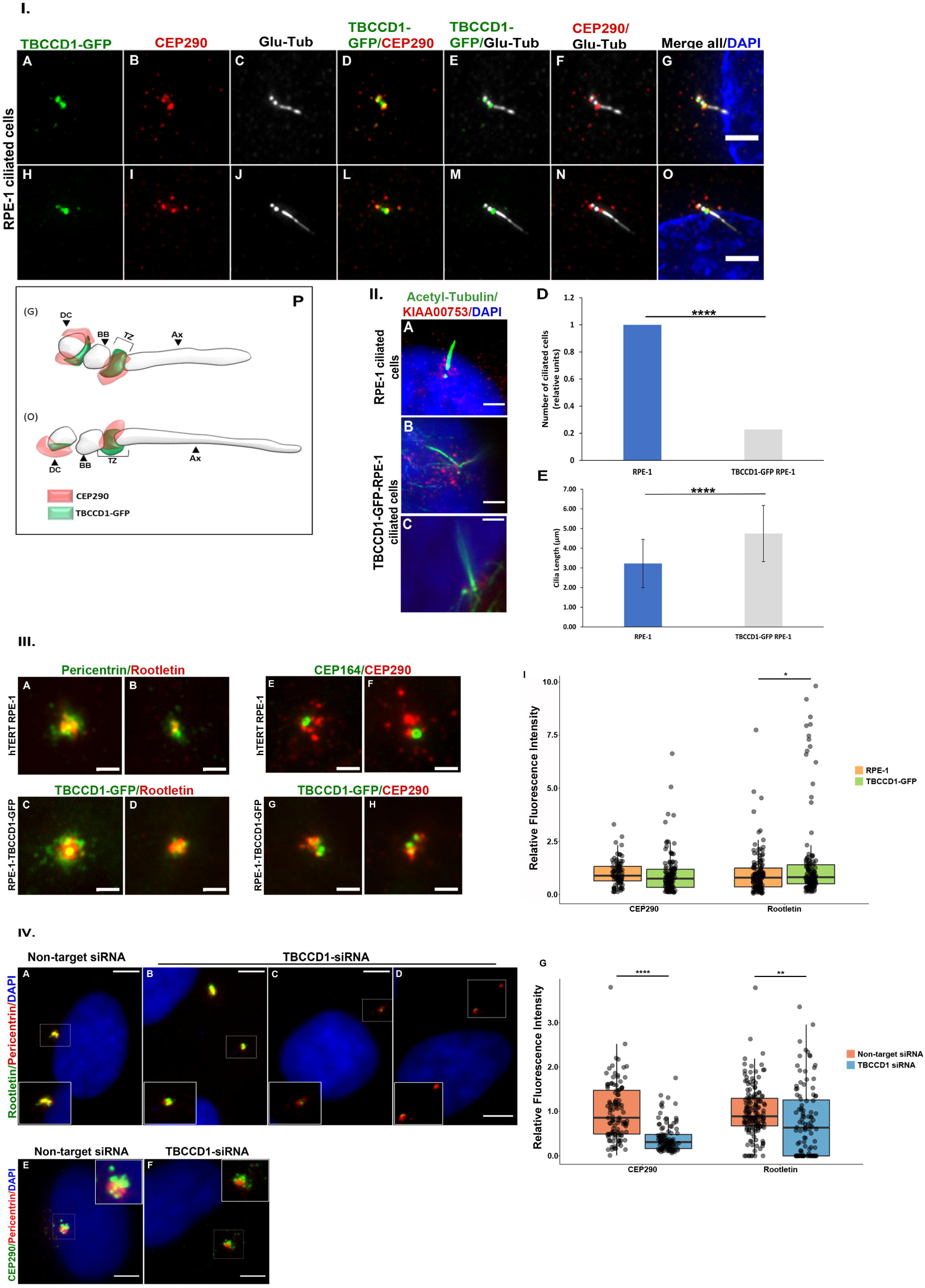
TBCCD1 recruits Rootletin and CEP290 to the centrosome and localizes at the distal end of the BB of human primary cilium. **I.** Immunofluorescence microscopy images of ciliated RPE-1 TBCCD1-GFP cells. **A to O** - TBCCD1-GFP (green), CEP290 (red) is used as a marker of the cilium transition zone, glutamylated tubulin (white) is used as a cilium/BB marker, and DNA is stained with DAPI (blue). Scale bars = 3 μm. **P** - Schematic representations of the BB localization of TBCCD1 and the transition zone marker CEP290. DC – daughter centriole; BB – basal body; TZ – transition zone; Ax – axoneme. **II.** Immunofluorescence microscopy images of ciliated RPE-1 and RPE-1 overexpressing TBCCD1-GFP cells. **A** – Acetylated tubulin (green) was used as a cilium axoneme marker, KIAA0753 (red), and DAPI shows DNA stained in blue. Scale bars = 1 μm. **B and C** –Acetylated tubulin (green) is used as a cilium marker, KIAA0753 (red), and DAPI shows DNA stained in blue. Scale bars = 1 μm. **D** – Percentage of RPE-1 and RPE-1 TBCCD1-GFP cells with a cilium in relative units. Three independent experiments were performed with n > 250 cells for each condition. **E** – Average cilium length in RPE-1 and RPE-1 TBCCD1-GFP cells. Cilium length was measured using Fiji – ImageJ. Error bars represent standard deviation. Significant differences to the corresponding control are indicated by **** (t-Student test; p < 0.0001). **III.** Immunofluorescence microscopy images of Rootletin and CEP290 in RPE-1 and RPE-1 TBCCD1-GFP cells. **A and B** –RPE-1 stained for Rootletin (red) and Pericentrin (green) as a centrosome marker. **C and D** – Impact of TBCCD1-GFP (green) overexpression in Rootletin (red). Scale bars = 1 μm. **E and F** –RPE-1 stained for CEP290 (red) and CEP164 (green) as a mother centriole marker. **G and H** – Impact of TBCCD1-GFP (green) overexpression in CEP290 (red). Scale bars = 1 μm. **I** - Relative fluorescence intensity for centriolar proteins CEP290 and Rootletin in RPE-1 cells constitutively expressing TBCCD1-GFP compared to RPE-1 cells. Fluorescence intensity was measured using Cell Profiler v4.2.1. Three independent experiments were performed with n > 130 cells for each condition. Grey dots represent individual measurements. Boxplot represented in colors. Significant differences to the corresponding control are indicated by * (t-Student test; p < 0.05). **IV** - Immunofluorescence microscopy localization of CEP290 and Rootletin in RPE-1 cells subjected to non-target siRNA and RPE-1 cells depleted of TBCCD1 by siRNAs. **A-D** - Impact of TBCCD1 depletion by siRNAs in Rootletin (green). Pericentrin (red) was used as a centrosome marker, and DAPI stains DNA in blue. Rootletin levels were heterogeneous among the mislocated centrosomes, with some showing that Rootletin was almost completely lost (D). **E-F** – Impact of TBCCD1 depletion by siRNAs in CEP290 (green). Pericentrin (red) was used as a centrosome marker, and DAPI stains DNA in blue. CEP290 levels were decreased by TBCCD1 depletion. Images are representative of three independent experiments. Inserts represent zoom images of centrosomes surrounded by punctuated white squares. Scale bars = 5 μm. **G** - Relative fluorescence intensity of centriolar proteins CEP290 and Rootletin in TBCCD1 depletion background in RPE-1 cells using specific siRNAs in comparison to RPE-1 cells treated with a non-target siRNA Fluorescence intensity was measured using Cell Profiler v4.2.1. Grey dots represent individual measurements. Boxplot represented in colors. Significant differences to the corresponding control are indicated by ** (t-Student test; p < 0.01) and **** (t-Student test; p < 0.0001).

Considering that, in paramecia, low levels of TBCCD1 caused aberrant number, length, and antero-posterior orientation of BBs’ accessory structures (striated fibers and microtubular rootlets), we then asked whether altered levels of TBCCD1 (by depletion and overexpression) affected Rootletin levels at the centrosome. Rootletin, which plays a role in centrosome cohesion in cycling cells (Bahe et al., 2005), is a structural component of the ciliary striated rootlet, a cytoskeleton-like structure originating from the BB and extending proximally toward the nucleus (Chen et al., 2015; Yang et al., 2002). Increased levels of TBCCD1 caused a 50% increase in Rootletin levels (n=130) (Figure 8-III, panels A-D and I). On the other hand, TBCCD1 depletion led to a 23 % decrease in Rootletin levels (n=107 centrosomes) (Figure 8-IV panels A-D and G). In these latter experiments, the decrease of Rootletin levels was heterogeneous among the affected centrosomes, with some almost devoid of Rootletin (29.9 %) (see Figure 8-IV, panels B-D and G). Interestingly, in centrosomes where Rootletin was almost completely lost, only 34.4 % showed centriole splitting (see Figure 8-IV, panel D). The loss of Rootletin from the centrosome caused by TBCCD1 RNAi seems not to be due to the loss of Rootletin interactor C-NAP1 since its levels were not affected by TBCCD1 depletion (Figure 5 I, panels A-B and P). These data suggest that multiple factors are required to maintain centrosome cohesion, including TBCCD1.

Together, our results show that TBCCD1 localized at the distal end of the BB, extending to the cilia transition zone in RPE-1 cells, and showed that cilia assembly requires strictly regulated levels of TBCCD1. Changes in TBCCD1 levels differentially impacted components of the mother centriole that are required for its successful conversion into a BB, as well as for its anchoring and assembly of primary cilia. Indeed, deregulated levels of TBCCD1 affected cilia assembly efficiency and length, probably by affecting the levels of some mother centriole appendage proteins (CEP164, Centriolin/CEP110, Ninein, and CEP170), the distal protein CEP290 as well as CEP350. For example, overexpression of TBCCD1 led to decreased levels of the DA protein CEP164 (Figure 5 III), which is required for BB anchoring and cilia assembly (Reed et al., 2022). In contrast, it leads to an increase in levels of CEP350 (see Figure 5-III), a protein that has also been shown to be involved in IFT complexes’ ciliary entrance (Kanie et al., 2017) which are involved in cilia length regulation (Keeling et al., 2016). The decrease of Rootletin levels, in response to either low or high levels of TBCCD1, highlighted the importance of TBCCD1 for centrosome cohesion in cycling cells and, most probably, for centrosome positioning (Potter et al., 2017), as well as for the assembly of the striated rootlets (bundles of MTs) required for primary cilia assembly and BB anchoring.

### 6 The proximity interaction network of TBCCD1

To gain further insights into the molecular mechanisms TBCCD1 is involved in, we performed a BioID analysis to establish its proximity interaction network. TBCCD1 was cloned in the pcDNA5 FRT/TO-BirA* vector in fusion with a FLAG-BirA* tag at the N-terminal end. This construct was used to transfect Flp-In TRex HEK293 cells and generate the stable/inducible cell line FLAG-BirA*-TBCCD1. Similarly, and in parallel, we used the empty pcDNA5 FRT/TO-BirA* vector to generate a negative control cell line expression FLAG-BirA* only. The expression of the TBCCD1 protein was induced with optimized concentrations of tetracycline to avoid high levels of TBCCD1 expression (see Figure S6). The centrosomal localization of TBCCD1 and the *in vivo* biotinylation at the centrosome were confirmed by immunofluorescence (IF) microscopy using γ-tubulin as a centrosomal marker and streptavidin labelling, respectively (Figure S5). This analysis confirmed the localization of TBCCD1 and the occurrence of biotinylation at the centrosome. Experiments were performed to show that the synthesis of TBCCD1 was induced by tetracycline to reveal the protein biotinylation pattern and the specific immunoprecipitated pattern of biotinylated proteins from FLAG-BirA*-TBCCD1 cells and from parental Flp-In-Trex and FLAG-BirA* cells (see Figure S7 for detailed descriptions). Considering the obtained results, we proceeded with the BioID analysis using FLAG-BirA* and FLAG-BirA*-TBCCD1 cells accordingly to material and methods.

BioID analysis identified 82 high-confidence TBCCD1 proximity interactors (Figure 9A, 9B, and Supplementary Table S1). We used Cytoscape to visually organize the TBCCD1 proximity interaction network (see material and methods and Figure 9B), which revealed a core of seven proteins (CEP162, KIAA0753/Moonraker/OFIP, MED4, OFD1, PCM1, RPGRIP1L, SSX2IP) (Figure 9B) connected with four protein clusters comprising most of TBCCD1’s interactome. A Gene Ontology (GO) enrichment analysis revealed a TBCCD1 interactome enriched in centrosomal, cytoskeleton, and cilium proteins (Figure S9 for details, see Supplementary Table S2). The interactome of TBCCD1 showed the highest enrichment for biological processes and/or pathways related to the centrosome, cytoskeleton, and cilia, including protein localization to the centrosome and MT cytoskeleton, centrosome organization and duplication, namely centriole assembly and replication, and in cilium assembly and ciliary BB-plasma membrane docking (Figure 9D; for details see Supplementary Table S3). This same analysis showed that five of the seven core proteins were associated with GO terms related to cilium assembly and organization (CEP162, OFD1, PCM1, RPGRIP1L, SSX2IP), and there was a high enrichment of centriolar satellites proteins (KIAA0753, OFD1, PCM1, SSX2IP) and ciliary BB proteins (OFD1, PCM1, RPGRIP1L, SSX2IP). Of the 4 clusters obtained in Cytoscape (Figure 9B), we defined clusters of proteins associated with: (i) centriole and cilium assembly (blue); ii) protein localization to centrosome and MT cytoskeleton (yellow); iii) cytoskeleton organization and cell division (red)and iv) Wnt signalling and cell polarity (green). Also, 28 proteins in the TBCCD1 network were not clustered when undergoing GO analysis for biological processes.

**Figure 9.**
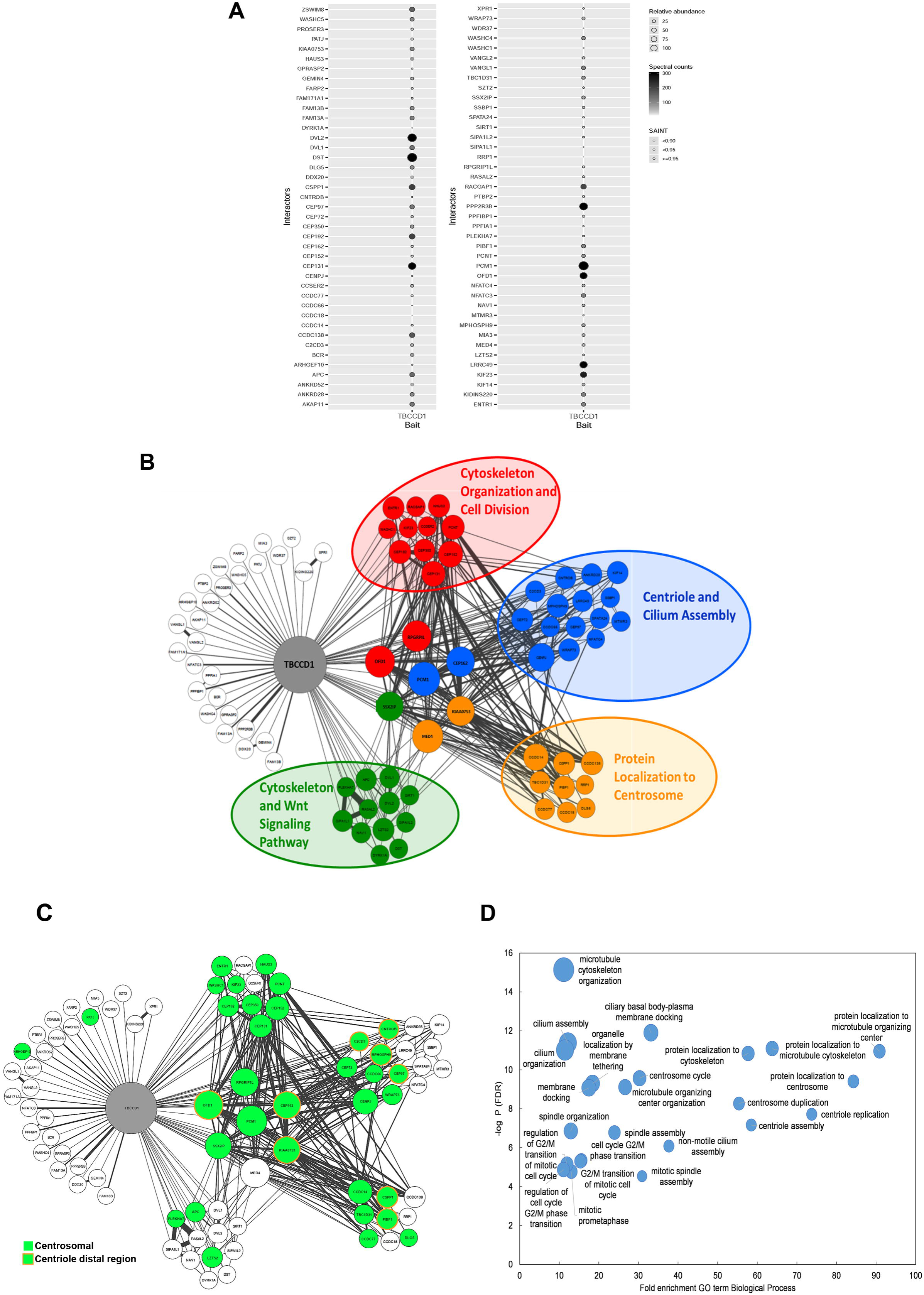
The vast interaction landscape of TBCCD1 is associated with the cytoskeleton, centrosome, and cilium. **A-** Mass spectrometry Dot Plot view of bait-prey interactions. The circle color (grey–black gradient) indicates the total number of spectral counts detected for each prey protein. Circle size indicates the relative abundance of prey protein in each BioID analysis, and circle color lines indicate the SAINT value (significance analysis of interactome; Teo et al., 2014); **B**- Protein physical interactions for each TBCCD1 BioID interactor were retrieved from public interactome databases using the PSICQUIC R package. The direct interactors subnetwork was visualized with the Cytoscape software (version 3.8.2; Shannon et al., 2003); **C** - Protein physical interactions for each TBCCD1 BioID interactor where centrosomal proteins (green) are highlighted. Proteins localized at the distal end of centrioles were signaled (orange circle) based on current literature (C2CD3- Tsai et al., 2019; Centrobin (CNTROB)- Gallaud et al., 2020; MPHOSPH9- Huang et al., 2018; CEP97- Spektor et al., 2007; CSPP1- Frikstad et al., 2019; CEP90- Le Borgne et al., 2022; OFD1- Singla et al., 2010; CEP162- Wang et al., 2013; KIAA0753-Kumar et al., 2021). **D** - GO enrichment analysis Biological Processes performed using the Gene Ontology Resource (http://geneontology.org/). X-axis presents the fold enrichment for the Biological Processes GO term; Y-axis presents the -log P for the False Discovery Rate (FDR). Circle size is proportional to the number of proteins of the TBCCD1 interactome associated with each GO term. Only terms with a Fold enrichment > 10 and a -log P (FDR) > 5 are presented.

In figure 9 C, we highlight the proteins that, by GO analysis, have a centrosomal localization and those known from the literature to have a distal centriolar localization as TBCCD1 (see figure 9 C legend for references). Indeed, 44% of the proteins that are members of the TBCCD1 proximity network are related to the centrosome, and 25% of these are described to localize at the distal end of one of the centrioles or both.

The literature data regarding the localization and functions of these TBCCD1 proximity interactors, combined with our results, clearly indicates that at the mother centriole distal end, TBCCD1 is in an environment composed of proteins that most probably regulate the remodeling of centriolar appendages occurring during the complex process of mother centriole to BB conversion, and its anchoring to the membrane. In conclusion, the proximity interactome at the distal end of the mother centriole supports that TBCCD1 participates in appendages’ maintenance/assembly and BB anchoring during ciliogenesis (Figures 6 and 7).

## DISCUSSION

Human TBCCD1 is a centrosomal protein related to the GTPase activating proteins tubulin cofactor TBCC and RP2 but does not seem to have this activity (Gonçalves et al. 2010). This protein is involved in centrosome positioning, cilia assembly, cell division, MT organization, and cytoplasm architecture (Godfroy et al., 2017; André et al., 2013; Gonçalves et al., 2010; Feldman et al., 2009). In this study, we identified TBCCD1 as an essential protein for the maintenance of the DA and SDA of the mother centriole. We also provide evidence that in motile cilia, TBCCD1 is required for correct BB positioning, orientation, and anchoring at the plasma membrane, as well as BB-accessory structures. Ultimately, our data link TBCCD1 cellular functions to the architecture of the cytoplasm space by its role in organizing the MT aster in interphase cells and to the remodeling of the distal region of the mother centriole and its anchoring to the membrane during cilia assembly.

### TBCCD1 levels differentially affect distal and subdistal appendages of the mother centriole, impacting MT anchoring to the centrosome

Here we show for the first time, using widefield, super-resolution microscopy alone or combined with expansion microscopy, that TBCCD1 decorates the two centrioles at the proximal and distal ends, being more abundant in the mother centriole. At the most distal end of the mother centriole, we observed a TBCCD1 complex (Figures 3 and 4) composed of a discrete ring that localizes at the level of the DA protein CEP164 connected to a conical structure inside the centriole barrel. This conic structure ends at the level of SDA proteins Centriolin/CEP110, Ninein, and CEP170 (Figures 3 and 4). Globally, this structure resembles an upside-down umbrella where the rods link the most distal region’s discrete ring to the conical structure’s vertex inserted inside the mother centriole’s lumen. This complex structure was always absent from the daughter centriole distal region, even though a small ring/dot of TBCCD1 was observed in some of these centrioles (Figures 3 and 4).

The mother centriole TBCCD1 distal complex structure establishes a spatial connection between the centriole region of the DA and that of the SDA. The existence of potential DA-SDA structural connections has recently been suggested by Chong et al. (2020), who showed that the structural arrangement of SDA is dependent on that of DA. For example, Ninein positioning at SDA is affected by DA structure in mutant cells for the DA protein CEP83 (Chong et al., 2020). The architecture of the TBCCD1 structure spanning from the DA to the SDA region fits well in a model where the crosstalk between these two structures can also be mediated by centriolar proteins that are not necessarily structural components of the appendages. Supporting this idea is the fact that TBCCD1 does not seem to be a component of SDA, as we showed by combined information from IF microscopy and the TBCCD1 proximity interaction network (Figure 9 and S9). However, its depletion causes the loss of CEP170 from the tip of SDA, accompanied by the decrease of centriolar levels of Centriolin/CEP110 and Ninein (Figure 5). Similarly to TBCCD1, the CEP350 protein, a member of the TBCCD1 proximity interaction network (Figure 9), is required for DA and SDA assembly without being a component of those structures (Karasu et al., 2022). CEP350, which localizes between the FOP protein and the centriole’s wall, is a key protein for regulating the length, stability, and assembly of the distal region of the centriole (Karasu et al., 2022). CEP350 also collaborates with EB1 and FOP in MT anchoring to the centrosome (Yan et al., 2006). Additionally, CEP350 stabilizes MT networks and the MTs of procentrioles (Clech, 2008; Hoppeler-Lebel et al., 2007) and forms a complex with FOP and CEP19 required for ciliogenesis (Mojarad et al., 2017). Interestingly, CEP350 levels are dramatically affected either by depletion or overexpression of TBCCD1 (Figure 5). In conclusion, although TBCCD1 not seemingly being a SDA component, it is required to stabilize Centriolin/CEP110 and the Ninein/CEP170 duo, which are involved in MT anchoring to the SDA (Delgehyr et al., 2005; Guarguaglini et al., 2005; Mogensen et al. 2000). Together these results provide a possible explanation for why centrosomes lost MTs in TBCCD1-depleted cells (Figure 1). On the other hand, we also observed that TBCCD1 depletion does not affect the DA protein CEP164 (Figure 5), which is required for DA assembly (Le Borgne et al., 2022; Kumar et al., 2021), showing that low levels of TBCCD1 probably do not compromise the assembly of DA. However, our studies showed that TBCCD1 overexpression led to a decrease of the levels of the DA protein CEP164, whereas it led to increased centriolar levels of the SDA proteins Centriolin/CEP110 and Ninein (Figure 5). These data suggest that the correct organization of the distal region of the mother centriole only occurs within a narrow range of TBCCD1 concentrations and that outside this optimal TBCCD1 concentration range, the DA and SDA are differentially compromised, impacting the cellular roles associated with these structures like, *e.g.,* cilia assembly.

The identity of the proximity distal centrosomal interactors of TBCCD1 (Figure 9), *e.g.*, C2CD3, OFD1, KIAA0753, and CEP90/PIBF1, also supports TBCCD1 as a player at the regulation of DA and SDA assembly/maintenance. Recently, it was shown that this group of proteins presents a unique molecular configuration at the base of the DA (Chang et al., 2023). In fact, most of these proteins seem to localize outside the centriole barrel at the distal end of one or both centrioles, with the probable exception of C2CD3 that, instead, seems to localize at the mother centriole’s distal end close to the centriole wall inside the barrel (Chang et al., 2023). In this group, OFD1 seems to play a pivotal role (see figure S9), showing an interaction with most of the proteins (C2CD3, KIAA0753, and CEP90) and forming a ternary complex with KIAA0753 and FOPNL/FOR20, which is present at the centrosome/centriole and pericentriolar satellites in human cells, and is required for ciliogenesis (Chevrier et al. 2016). KIAA0753 and OFD1 are involved in centriole length control and, together with CEP90, are required for DA assembly and cilia biogenesis (Kumar et al., 2021). OFD1 and C2CD3 are also involved in centriole length regulation and are required to recruit the distal appendage proteins CEP164, CEP83/CCDC41, CEP89, FBF1, and SCLT1 to the mother centriole (Ye et al., 2014; Tang et al., 2013; Singla et al., 2010). Moreover, C2CD3, OFD1, and Talpid3 differentially regulate the assembly of SDA, the CEP350/FOP/CEP19 module, centriolar satellites, and actin networks (Wang et al., 2018). This clearly supports the view that TBCCD1 localizes at the distal end of the mother centriole (see also Figures 3 and 4) in the neighborhood of a group of proteins involved in DA and SDA assembly (*e.g.,* OFD1), playing an essential role in cilia biogenesis.

Despite all the evidence that TBCCD1 plays a role in the distal region of the mother centriole related to appendage structure maintenance, the question remains of how TBCCD1 affects these structures without being their component. We put forward the hypothesis that at the mother centriole, the region defined by the SDA is continuously subjected to mechanical forces generated by the pushing and pulling forces of the MTs anchored at the tips of these structures. It is plausible that TBCCD1 is required to maintain the centriole mechanical properties to cope with the forces resulting from MTs anchoring/dynamics. Then, low levels of TBCCD1 will alter the mechanical properties of the centriole, weakening the stability of the SDA structure predominantly at the region where the forces are applied, *i.e*., the most distal part of the structure (far from the centriole barrel), where the module Ninein/CEP170 plays a role in MT nucleation and aster anchoring (Uzbekov and Alieva, 2018; Delgehyr et al., 2005; Guarguaglini et al., 2005; Dammermann and Merdes, 2002; Mogensen et al., 2000; Quintyne et al., 1999). A direct consequence of the structural disruption of the most distal region of the SDA would be a random release of MTs with a concomitant asymmetry of the MT aster, which was observed in RPE-1 TBCCD1 knockdown cells (Figure 1). Also, a direct outcome of the loss of MTs from SDA and their radial organization will probably be a perturbation of the balance of pulling/pushing forces exerted by the MTs, driving the centrosome from the centroid to the periphery of the cell (Odell et al., 2019; Tanimoto et al., 2018; Letort et al., 2016; Wu et al., 2011; Zhu et al., 2010; Kimura and Onami, 2010; Dogterom et al., 2005; Théry et al., 2006; Burakov et al., 2003), a phenotype observed when TBCCD1 is depleted (Gonçalves et al., 2010). On the contrary, increased levels of TBCCD1 increase the centrosome capacity to anchor MTs through the increased levels of Centriolin/CEP110, Ninein, and CEP350, perturbing the DA by reducing the levels of the DA protein CEP164 (Figure 5).

Moreover, in most of the analyzed centrioles, we found a TBCCD1 ring localizing at the proximal region of both mother and daughter centrioles, above the localization of C-NAP1, Ninein, and CEP170 (Figure 3). Consequently, TBCCD1 is a new member of the protein group with both distal and proximal localizations at the mother centriole, a group that includes Ninein, CEP170, Kif2A, p150glued, and CCDC68, CCDC120, and VFL3 (Pizon et al., 2020; Huang et al., 2017; Mazo et al., 2016). Interestingly, all these proteins have been associated with the SDA and have a role in MT anchoring, organization, and stability (Pizon et al., 2020; Huang et al., 2017; Mazo et al., 2016; Desai et al., 1999). At the proximal region of the centrioles, TBCCD1 knockdown also caused the loss of CEP170, which was accompanied by a decrease of the centrosomal levels of Rootletin and without significant changes in C-NAP1 levels (Figures 5 and 8). The loss of Rootletin is coherent with our previous observation that in RPE-1 TBCCD1-depleted cells, the dynamic movement between both centrioles in interphase cells was reduced (see supplementary movie 2 in Gonçalves et al., 2010). Rootletin has been implicated in centrosome cohesion by forming centriole-associated fibers that directly interact with C-NAP1 at the proximal end of centrioles (Vlijm et al., 2018; Yang et al., 2006; Bahe et al., 2005). C-NAP1 is also required for Ninein/CEP170 recruitment to this region (Mazo et al., 2016). Thus, our observations suggest TBCCD1 as a new factor stabilizing the centriolar proximal Ninein/CEP170 duo and the centrosome cohesion factor Rootletin independently of C-NAP1. Similarly to TBCCD1, the protein VFL3 was also described as able to regulate centriole tethering and to affect Rootletin independently of C-NAP1 (Pizon et al., 2020). Interestingly, TBCCD1 mutant *Chlamydomonas* cells show defects in centriole linkage (Feldman et al., 2009). Strikingly, our data also show that the localization of TBCCD1 at the proximal centriole region is important during the biogenesis of new centrioles since the protein localizes in the sites where the procentrioles start to assemble, suggesting that TBCCD1 localization at the mother centriole is closely related with its maturation stage (Figure 4).

Collectively, our data mainly demonstrate that, independently of the structures where TBCCD1 localized, it was required for centriolar localization of the MT nucleation/anchoring proteins, Ninein and CEP170 at either the distal end of MC or at the proximal end of both centrioles. Coherent with this conclusion is the observation that aggregates of acentriolar TBCCD1 contain Ninein and CEP170 and can nucleate and organize radial MT networks (Figure 2). We do not have data supporting TBCCD1 binding to MTs (which was also not supported by its proximity interactome), except at anaphase midzone MTs, where TBCCD1 is recruited from the cytoplasm and later accumulates at the midbody (Figure S2 and movie S1). At the midzone, TBCCD1 decorates small segments of the anti-parallel array of MTs, most probably through its partners. For example, the protein CEP97, a component of the proximity interactome of TBCCD1, is a binding partner of the centrosomal +TIP CEP104/KIAA0562 (Jiang et al., 2012). During cell division, the dynamics of astral and midzone MTs must be distinctly regulated, with the midzone MT stabilization coordinated with actomyosin contraction and furrow ingression (Landino et al., 2016). The centrosomal and acentrosomal localization of TBCCD1 strongly suggests that this protein is required to maintain structures involved in MT anchoring and that cope with mechanical forces.

### TBCCD1 is essential for motile cilia BB anchoring and for building the BB appendages

Using *Paramecium* as a biological model, we observed that TBCCD1 is essential for correctly positioning, orienting, and anchoring motile cilia BBs to the cell surface. Most probably, this TBCCD1 role is related to its requirement for the correct assembly and maintenance of BB accessory structures, namely the striated rootlet and the MT rootlets (Figures 6 and 7). It is well documented that *Paramecium* BB appendages are essential during BB biogenesis by acting as a scaffold for the new BB being required for its positioning and anchoring to the cell surface (Iftode and Fleury-Aubusson, 2003). In the ciliate *Tetrahymena,* it has been shown that the final positioning of the BBs in the ciliary row, including correct spacing and orientation, is dependent on the relationship between striated rootlets and the organization of the cell’s cortex components in a ciliary force-dependent mechanism (Soh et al., 2020; Galati et al., 2014; Allen 1969). Indeed, striated rootlets are dynamic structures that can alter their length to link neighboring BBs to each other and the cell cortex responding to changes in ciliary forces (Soh et al., 2020).

Interestingly, in *Paramecium*, the orthologue of human VLF3/CCDC61 gene encoding a protein of SDA of mother centriole (Pizon et al., 2020), when knocked down (Bengueddach et al., 2017) displays phenotypes that resemble those of *Paramecium* TBCCD1 loss of function. Accordingly, in the VFL3-A knockdown, the organization of BB appendages is affected, and the BBs lose their circumferential polarity (Bengueddach et al., 2017). TBCCD1 shares the ability to regulate the assembly/maintenance of BB accessory structures with VLF3-A. The decrease in *Paramecium* swimming speed observed in TBCCD1-depleted cells is consistent with the improper positioning of BB in longitudinal rows relative to the cellular anterior-posterior axis due to defects in striated and microtubular rootlets. Indeed, mispositioned/misoriented BBs should affect the direction and coordination of ciliary beating (Hoops et al., 1984; Gibbons, 1981; Tamm et al., 1975).

The decreased ability of RPE1 cells to assemble primary cilia may also be related to deficiencies in the assembly of BB-associated structures. For example, Rootletin also organizes the ciliary rootlet, a cytoskeleton-like structure that emanates from the BB toward the cell nucleus and is required for vertebrate ciliary maintenance and function (Styczynska-Soczka and Jarman et al., 2015; Chen et al., 2015; Mohan et al., 2013;). The ciliary rootlet is analogous to the striated rootlet of *Paramecium* BBs (Chien et al., 2013) and is found throughout the eukaryote phylogenetic tree, although with different compositions. Our results (Figures 7 and 8; Gonçalves et al., 2010) support the idea that throughout evolution, Rootletin has evolved to substitute the components of lower eukaryotes’ striated rootlets, namely the striated fiber assemblins (Joukov and Nicolo, 2019). These structures probably evolved to transduce ciliary forces to BB positioning in motile cilia (Soh et al., 2020) and mechanosensing as in sensory neuron cilia (Styczynska-Soczka and Jarman, 2015; Chen et al., 2015). During evolution, it seems that TBCCD1 kept the role required to maintain these structures, independently of their composition and most probably by maintaining the structural stability and integrity of the accessory structures of these BB (Figure 8).

Human primary cilia also have basal feet, structures that are important to maintain the primary cilium submerged in the cytoplasm by linking the BB to the Golgi (Galati et al., 2016; Mazo et al., 2016). These structures seem to organize from the mother centriole SDA in coordination with their centrosomal proximal pool (Nguyen et al., 2020). Recent super-resolution data suggest that these structures have a modular architecture involved in BB anchoring, scaffolding, and MT organization, where Centriolin/CEP110, Ninein, and CEP170 play an important role (Nguyen et al., 2020). Taking into account the impact TBCCD1 depletion had in these SDA proteins and in the Ninein/CEP170 proximal pools (Figure 5), we envisage that TBCCD1 depletion might affect primary cilia basal feet, which could contribute to the lower ability of RPE-1 cells to assemble primary cilia (Gonçalves et al., 2010).

Our results also suggest that TBCCD1, in human cells, may have a role in the maturation of the distal region of the mother centriole during the conversion to BB. In fact, TBCCD1 depletion caused a decrease in the levels of CEP290 in RPE-1 cells, associated with a lower ability to assemble primary cilium (Figure 8 and Gonçalves et al., 2010). On the other hand, RPE-1 cells with high levels of TBCCD1 do not have altered CEP290 levels but also show a decrease in the efficiency to assemble primary cilia. Moreover, cilia tend to present longer axonemes in TBCCD1 overexpressing cells. This may be related to the fact that TBCCD1 overexpression in RPE-1 cells causes the decrease in CEP164 levels since this protein mediates the vesicles docking to the mother centriole, BB anchoring, and IFT transport during primary cilia assembly (Reed et al., 2022; Schmidt et al., 2012). Supporting this idea, the protein MPHOSPH9, a proximity partner of TBCCD1, is required for cilia formation by promoting the removal of the CP110-CEP97 complex from the distal end of the mother centriole (Huang et al., 2018). CEP162, also a TBCCD1 proximity partner, displays MT-binding activity and is tethered at the mother centriole distal ends promoting TZ assembly in primary cilia (Wang et al., 2013), whereas CSPP1 is involved in the regulation of axoneme length (Frikstad et al., 2019). The altered length of primary cilia assembled in the TBCCD1 overexpression background may also be related to alterations in the transition zone structure that do not involve CEP290. In fact, some transition zone proteins, e.g., RPGRIP1L/NPHP8 (a TBCCD1 proximity partner), have also been shown to be essential regulators of ciliary length (Wiegering et al., 2021; Liu et al., 2011; Patzke et al., 2010).

From our studies, we show that the TBCCD1 role is critical for microtubular structures that usually cope with forces generated by associated MTs like the SDA of the mother centriole, the proximal end of both centrioles, the spindle midzone/midbody, the BB distal end/cilia transition zone and the maintenance of accessory structures of BBs. In these structures, TBCCD1 plays an essential role in the organization and maintenance/recruitment of the proteins involved in the MT nucleation and anchorage, a role which may also occur at the cytoplasm where a cytoplasmic pool of this protein is observed (Figure S1). The vast proximity-interactome of TBCCD1 supports these functions (see Figure 9) and mainly explains the depletion phenotypes described in human cells (this paper and Gonçalves et al., 2010).

Finally, attention should be paid to the potential role of TBCCD1 in the development of ciliopathies. In fact, a screening using KEGG Mapper (Kanehisa and Sato, 2020) (Supplementary Table S3) reveals that nine proteins belonging to the TBCCD1 interactome are associated with ciliopathies or ciliopathies’ phenotypes such as primordial dwarfism (Shaheen et al., 2012), including Joubert’s syndrome (OFD1; RPGRIP1L; CSPP1; CEP90), orofacial syndrome (OFD1; C2CD3), COACH syndrome (RPGRIP1L), Meckel syndrome (RPGRIP1L; KIF14), Seckel syndrome (CENPJ; CEP152) and microcephalic osteodysplastic primordial dwarfism, type II (PCNT). Some of the proteins are associated with neurodevelopmental disorders, including primary microcephaly (CENPJ; CEP152; KIF14) and neurodevelopmental disorder with microcephaly (GEMIN4) or autosomal recessive mental retardation (WASHC4).

## ACKNOWLEDGMENTS

We thank Laurence Pelletier (Lunenfeld-Tanenbaum Research Institute, Sinai Health System, & Department of Molecular Genetics, University of Toronto, Toronto, Canada) for the kind gift of Flp-In T-REx 293 stable lines expressing FLAG-BirA*, to Erich Nigg (Biozentrum, University of Basel, Basel, Switzerland for antibodies against CEP164, CEP170, and C-NAP1 and to Andreas Merdes (Centre de Biologie Intégrative, Université Paul Sabatier/Centre National de la Recherche Scientifique, Toulouse, France) for providing the antibody against PCM1. We would also like to thank Carsten Janke for his generous gift of anti-poly-E antibodies. Finally, we are thankful to Mónica-Bettencout Dias for the use of the super-resolution microscope.

Centro de Química Estrutural is funded by Fundação para a Ciência e Tecnologia-projects PIDDAC-UIDB/00100/2020 and UIDP/00100/2020, as well as the Associate Laboratory Institute of Molecular Sciences: project LA/P/0056/2020. This work was also funded by Instituto Politécnico de Lisboa IPL/2016/TBCCentro_ESTeSL, IPL/2017/CILIOPAT/ESTeSL, IPL/2019/MOONOFCILI/ESTeSL and IPL/2021/ObeCil_ESTeSL. The present work has benefited from Imagerie-Gif core facility supported by I’Agence Nationale de la Recherche (ANR-11-EQPX-0029/Morphoscope, ANR-10-INBS-04/FranceBioImaging; ANR-11-IDEX-0003-02/ Saclay Plant Sciences). This work has been founded by “Basal body anchoring in ciliogenesis: structure-function analysis: ANR-15-CE11-0002-01” to AMT.

## MATERIALS AND METHODS

### 1 Cell lines, strains, and culture conditions

#### Human cells

293 Flp-In T-REx cells (a kind gift of Laurence Pelletier, Lunenfeld-Tanenbaum Research Institute Mount Sinai Hospital, Toronto, Canada) were grown in Dulbecco’s Modified Eagle’s Medium (DMEM) supplemented with 10% fetal bovine serum (FBS), GlutaMAX, zeocin (100 μg/ml) (Invitrogen, Paisley, UK) and blasticidin (3 μg/ml) (Thermo Scientific, Paisley, UK). Similarly, Flp-In T-REx 293 stable lines expressing FLAG-BirA* or FLAG-BirA* fusion TBCCD1 were grown as mentioned above after the addition of hygromycin B (200 μg/ml) (Invitrogen, Paisley, UK). The hTERT RPE-1 (RPE-1) and TBCCD1-GFP-hTERT RPE-1 (TBCCD1-GFP) cells were grown in DMEM/F12 supplemented with 10% FBS, GlutaMAX and sodium bicarbonate 0.25%. All cell lines culture occurred in a 5% CO_2_ humidified atmosphere at 37°C. The generation of a stable TBCCD1-GFP RPE-1 cell line was previously described by Cardoso et al. (2014).

To induce primary cilia assembly, RPE-1 and RPE-1-TBCCD1-GFP cells were seeded at a 2.5-fold higher density than 2 x 10^4^ cells in a 24-well plate, serum-starved for 24 h and then processed for immunofluorescence.

#### *Paramecium* strains

In RNAi experiments using the protozoa ciliate *Paramecium tetraurelia* wild-type strain, the Stock d4-2 was used. In transformation assays, it is required that cells do not discharge trichocysts and, therefore, the mutant cells nd7-1 were used (Skouri and Cohen, 1997). The cells were grown at 27 ° C in a wheat grass powder infusion (Bio Herbe de Blé L’arbre de vie, Luçay Le Male, France) bacterized with *Klebsiella pneumoniae* and supplemented with 0.8 µg/ml β-sitosterol according to standard procedures (Sonneborn, 1970).

### 2 MT depolymerization and polymerization assays

To depolymerize MTs, RPE-1 and TBCCD1-GFP cells were incubated with 30 µM nocodazole (Sigma-Aldrich) in dimethyl sulfoxide (DMSO) (Sigma, St. Louis, EUA), and the cell plates were put on ice during the treatment for 30 min. Subsequently, cells were carefully washed twice with culture medium for MT polymerization recovery and placed in fresh medium at 37°C for different times before they were fixed and processed for immunofluorescence.

In the rescue assays of the siRNA TBCCD1 phenotypes, 48 h post-transfection with TBCCD1 siRNAs, the culture media of RPE-1 cells was supplemented for 24 h with 0.1 and 0.2 nM Paclitaxel (Taxol; Sigma, St. Louis, EUA) in DMSO. The same DMSO volume was used as a control in both nocodazole- and Paclitaxel-treated cells.

### 3 Generation and characterization of FLAG-BirA*TBCCD1 and FLAG-BirA* cell lines for BioID

The coding sequence of the human *tbccd1* gene was amplified by PCR using the primers TBCCD1Forward 5’-TTGGCGCGCCTATGGATCAGTCCAGAGTTCTC-3’and TBCCD1Reverse 5’-ATAAGAATGCGGCCGCTTATCCAGCTGCTTGTTTGGA-3’ from the TBCCD1-pIC111 recombinant expression vector described in Gonçalves et al., (2010). The PCR product was cloned into the expression vector pcDNA5 Flag/FRT/TO-BirA* (a kind gift of Laurence Pelletier, Lunenfeld-Tanenbaum Research Institute Mount Sinai Hospital, Toronto, Canada) using the restriction enzymes NotI e AscI and the obtained recombinant vector was sequenced to confirm the absence of TBCCD1 coding sequence alterations.

For plasmid transfections of the human cell lines, Lipofectamine-2000 or Lipofectamine-3000 (Invitrogen, Paisley, UK) was used according to the manufacturer’s instructions. The HEK293T Flp-In/T-Rex cells were co-transfected with pOG44 (Flp-recombinase expression vector), and the plasmid containing only the coding sequence for FLAG-BirA* TBCCD1 or for FLAG-BirA*. After transfection, cells were selected with blasticidin and hygromycin B, as described above.

To test TBCCD1 localization, protein biotinylation and evaluate overexpression artifacts, Flag-BirA* TBCCD1 and Flag-BirA* cell lines were incubated for different times in complete media with (a) 0.5 μg/mL or 1 μg/mL tetracycline (SIGMA) and 50 μM biotin (Sigma-Aldrich), or (b) only) 0.5 μg/mL or 1 μg/mL tetracycline, or (c) in the absence of both tetracycline and biotin and then processed for immunofluorescence microscopy or protein analysis by SDS-PAGE followed by Western-blot.

### 4 BioID sample preparation and Mass spectrometry

To determine the TBCCD1 interacting network, we used the BioID method (Roux et al., 2012) and the protocol described in Gupta et al., 2015. The SAINT software (Significance Analysis of INTeractome) was used to select the most likely interacting proteins from those found by mass spectrometry analysis. The SAINT software assigns a degree of confidence that varies between 0 and 1 (SAINT value) for all proteins found in the analysis, being selected those that have a degree of confidence greater than 0.8 (Choi et al., 2011). The results were obtained from 2 biological replicates and two technical replicates (*i.e*., each replicate was analyzed twice).

### 5 Data analysis and visualization tools of the TBCCD1 proximity interaction network

Protein physical interactions for each TBCCD1 BioID interactor were retrieved from public interactome databases using the PSICQUIC R package. The selected database providers where: BioGrid, IntAct, IMEx, mentha, MatrixDB, Reactome, and BindingDB.

The igraph R package was used to analyze the resulting protein interaction network. Neighbor similarity between nodes was computed with Jaccard coefficients. The Jaccard similarity coefficient of two nodes is the number of common neighbors divided by the number of nodes that are neighbors of at least one of the two nodes being considered. A sub-network containing only TBCCD1 and its direct BioID interactors was extracted for visualization and further analysis. Proteins in this subnetwork were grouped into clusters that maximized network modularity with the function “cluster_fast_greedy”. Large cliques were also detected with the function “largest_cliques”. The direct interactors subnetwork was visualized with the Cytoscape software (version 3.8.2) (Shannon et al., 2003). The yFiles organic layout was first applied and later manually edited to a) group proteins within the same cluster and b) group proteins that are part of the largest cliques.

TBCCD1 proximity interaction network proteins classification was performed using the PANTHER classification system (Mi et al., 2019). Gene Ontology (GO) (Ashburner et al., 2000) enrichment analysis was performed using the Panther Gene Ontology resource (http://geneontology.org/) (Mi et al., 2019). P values were obtained using Fisher’s exact test and corrected for false discovery rate (FDR). KEGG enrichment analysis was performed using the STRING database and KEGG Mapper (Kanehisa and Sato (2020) to see if the TBCCD1 interactome proteins were associated with diseases.

### 6 Protein extracts, immunoprecipitation, and Western blot

In case of immunoprecipitations, Flag-BirA* Tbccd1 and Flag-BirA* cells were induced by incubating them with tetracycline (1 µg/mL) for 24 h, were washed, harvested, and frozen at −80°C for 5-10 min or lysed immediately. For total protein extracts, the thawed pellets were resuspended in lysis buffer (50 mM HEPES-KOH; pH 8, 100 mM NaCl, 5 mM EDTA, 0.2 % (v/v) NP-40, 250 mM sucrose; protease and protease inhibitor cocktail (Thermo Scientific) for 1 h at 4° C with gentle rotation. Next, nucleases (Pierce™ Universal Nuclease for Cell Lysis) or benzonase nuclease (MilliporeSigma) were added to extracts cleared by sonication. Then lysates were centrifuged at 14,000 *g* for 40 min at 4° C, and the supernatant was collected. The cytoplasmic and nuclear extracts were prepared as described in Gonçalves et al. (2010).

A fraction of the protein extracts obtained was saved to be used later as (Inputs), and the remaining was incubated overnight at 4°C with Dynabeads M-280 Streptavidin (Invitrogen, Paisley, UK) with gentle agitation. After the incubation, the beads were pelleted and washed several times with Tween 0.01% (v/v) 1x (4.3 mM Na_2_HPO_4_, 1.4 mM KH_2_PO_4_, 137 mM NaCl and 2.0 mM KCl, pH 7.4) PBS and twice with 1x PBS.

For total protein extracts to be directly analyzed by Western blot, the cells were collected, lysed in Laemmli buffer, and treated with nucleases. Immunoprecipitated and total protein extracts were analyzed by SDS-PAGE 10% (w/v) gels, and proteins were detected with silver staining *Pierce Silver Stain* (Thermo Scientific, Paisley, UK) accordingly to manufacturer’s instructions. For immunodetection, the proteins were transferred to nitrocellulose. Blots were blocked at room temperature for 1 h in PBS (4.3 mM Na_2_HPO_4_, 1.4 mM KH_2_PO_4_, 137 mM NaCl and 2.0 mM KCl, pH 7.4) containing 5% (w/v) fat-free milk powder before incubation with primary antibody. The primary antibodies and dilutions used were: anti-GFP (Roche, dil. 1:1000); anti-α-tubulin (Sigma, clone DM1A; dil. 1:1000); Anti-Flag M2 (Sigma, dil. 1:1000), and anti-TBCCD1 (Sigma, dil. 1:250). Biotinylated proteins were identified using HRP-conjugated streptavidin (Invitrogen, 1:10000). After extensive washing in PBS containing 0.1% (v/v) Tween-20, the blots were incubated with secondary-antibody-peroxidase conjugates (Jackson ImmunoResearch and Zymed) at 1:2000 dilutions, washed extensively with 1x PBS and then developed by enhanced chemiluminescence (ECL, GE Healthcare).

### 7 TBCCD1 genes cloning and *Paramecium* transformation

*Paramecium* constitutive cell lines expressing ciliate GFP-TBCCD1 gene (ParameciumDB GSPATP00004357001) were constructed by transforming the ciliate cells with the pPXV-GFP vector plasmid (kindly provided by Jean Cohen) where the TBCCD1 coding region, amplified from genomic DNA by PCR, was cloned under the constitutive regulators of the calmodulin gene, and downstream of the GFP-coding region (pPXV-GFP-TBCCD1).

Then, Nd7-1 mutant cells were transformed by microinjection in the macronucleus with a mixture of the linearized pPXV-GFP-TBCCD1 plasmid (at 5 μg/μl) and the plasmid allowing the expression of the *ND7* wild-type gene to complement the Nd7 mutation (Skouri and Cohen 1997). Microinjection was made under an inverted Nikon phase-contrast microscope using a Narishige micromanipulation device and an Eppendorf air pressure microinjector. The transformation was assessed by screening the ability of *Paramecium* to discharge trichocysts. The positive cells were then analyzed for GFP-TBCCD1 localization. Clones expressing the fusion protein and showing a growth rate similar to untransformed Paramecia were chosen for further analyses.

### 8 RNA interference

#### Human cells- TBCCD1 siRNAs

The RPE-1 cells (1 x 10^4^ cells seeded in 24-well plates) were transfected with 100 nM of a pool of four siRNAs directed to TBCCD1 obtained from Dharmacon (ON-TARGETplus Duplex; Lafayette, CO, USA) and Ambion (Silencer Select siRNAs; Austin, TX, USA) as described in Gonçalves et al. (2010), using Lipofectamine 3000 (Invitrogen, Paisley, UK) according to the manufacturer’s instructions. After 48 h post-transfection, RPE-1 cells were subjected to a new boost of siRNAs pool, and at 72 h post-transfection, cells were fixed and processed for IF. The Silencer Select Negative Control 2 siRNA (Ambion) was used as a negative control.

#### Gene silencing in Paramecium *cells by feeding*

*TBCCD1 depletion by RNAi*- The RNAi experiments *Paramecium* cells were performed accordingly to the feeding method described in Galvani and Sperling (2002). *Paramecium* cells were fed with HT115 DE3 *Escherichia coli* strain expressing specific T7Pol-driven double-stranded RNA for both *Paramecium* TBCCD1 genes (ParameciumDB GSPATP00004357001 and GSPATP00019048001). Paramecia were fed with these bacteria and refed daily with a fresh feeding medium at 27°C. Control cells were fed with HT115 bacteria carrying the L4440 vector containing the ND7 gene involved the exocytosis (Skouri and Cohen 1997). Paramecia transformed cells were transferred daily into fresh feeding medium. Phenotypes were analyzed after 48h of feeding. RNAi-off target effects were analyzed for each construct using the RNAi-off target tool available at ParameciumDB (Arnaiz et al., 2007). This approach originates distinct cell lines presenting homogeneous phenotypes. The effectiveness of RNAi was quantified by analyzing the decrease of BB fluorescence in transformed cell lines expressing the GFP-tagged transgene after silencing. In *Paramecium*, the phenotype of silenced cells is highly reproducible from one experiment to another and from one cell to another. In each experiment, paramecia were analyzed in at least 2 or 3 independent replicates.

To analyze swimming phenotypes, *Paramecium* control cells and those depleted in TBCCD1 were tracked for 10 s every 0.3 s using a Zeiss Steni 2000-C dissecting microscope with a Roper Coolsnap-CF camera and Metamorph software (Universal Imaging). Stacks were analyzed using the Manual tracking tool in ImageJ. For this, 4-8 *Paramecium* cells were collected and placed in 10 μL drops of bacterized BHB solution, where bacteria were removed, after their growth, by sterilization.

### 9 Immunofluorescence microscopy, super-resolution microscopy, and expansion microscopy

#### Human cells

For immunofluorescence microscopy, the cells were fixed with cold methanol (10 min at −20°C) and blocked for 30 min with 3% (w/v) bovine serum albumin (Calbiochem) in 1 x PBS at room temperature. Next, they were incubated for 1 h at room temperature with the primary antibodies (see table 2 for dilutions) diluted in the block solution. After incubation, cells were washed twice with 1 x PBS and once with 1 x PBS/ 0.1% (v/v) Tween-20 and then incubated for 1 h at room temperature with secondary antibodies (also diluted in block solution). For immunofluorescence using Super Resolution Microscopy, the cells were fixed with cold methanol (10 min at −20°C) and blocked for 30 min with 10% (v/v) fetal bovine serum (Gibco) in 1 x PBS at room temperature. Next, they were incubated for 1h at room temperature with the primary antibodies diluted in the block solution. After incubation, cells were washed three times with 1 x PBS and then incubated for 1h at room temperature with secondary antibodies (see table 2 for dilutions). Either for immunofluorescence microscopy or super-resolution microscopy, cells were then washed three times with 1 x PBS and incubated for 1-3 min at room temperature with 1 mg/mL DAPI and finally, coverslips were mounted onto glass slides by inverting them onto mounting solution Vectashield H-100 (Vector Laboratories, Inc). For the Expansion Microscopy, we used a modified protocol version of Gambarotto et al. (2019). 13 mm coverslips with cells were covered with a solution of 1.4 % formaldehyde and 2 % acrylamide in 1X PBS for 5h at 37° C. After the incubation, a drop of monomer solution (19 % sodium acrylate, 10 % acrylamide, 0.1 % N,N’-methylenbisacrylamide in 1 x PBS) with 0.2 % TEMED and 0.2 % ammonium persulfate (APS), was placed in parafilm inside a humid chamber and was rapidly covered with a coverslip with cells facing the solution. Gelation proceeded for 5 minutes on ice and then shifted to 37°C in the dark for 1 h. Then, coverslips with attached gels were transferred into a 6-well plate and incubated in 1 ml denaturation buffer (200 mM SDS, 200 mM NaCl, 50 mM Tris in water, pH 9) with agitation for 15 minutes. The gel was then transferred to an Eppendorf tube filled with denaturation buffer and incubated for 1 hour at 95° C. Gels were removed from the coverslips and placed into glass plates filled with ddH_2_O for expansion. Water was exchanged at least twice every 30 min and then incubated in 1X PBS for 15 minutes. The gels then proceeded to antibody incubation. In microtubule depolymerizing assays prior to methanol fixation, cells were rinsed with PBS1x, washed in Triton X-100 0.1% (v/v) in BRB80 (80 mM PIPES-KOH pH 6.8, 1mM MgCl_2_, 1 mM EGTA) and then fixed. This approach removes the soluble unpolymerized tubulin heterodimers, dramatically reducing background.

**Table 2.**
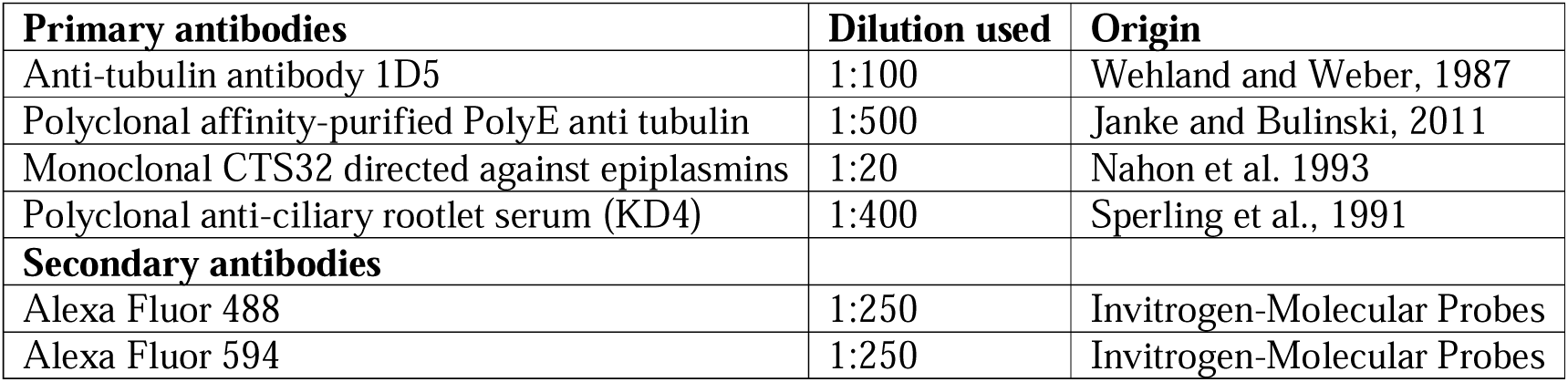
Antibodies and dilutions used in *Paramecium* cells.

#### Paramecium *cells*

In the cases of ciliate protozoa *Paramecium,* fixation, and immunofluorescence techniques were performed on cells in suspension. 50-100 cells were collected in the smallest volume possible and were permeabilized in 200 µl PHEM (Pipes 60 mM, Hepes 25 mM, EGTA 10 mM, MgCl_2_ 2 mM, adjusted to pH 6.9) with 1% Triton-X-100 (PHEM-Triton) for 30 secondes. Then cells were fixed for 10 min in 2% PHEM-PFA. Buffer was then aspirated, and cells were rinsed three times for 10 min in PBS-tween 0.1% with 3% BSA.

The human cells were analyzed using an Applied Precision DeltavisionCORE system, mounted on an Olympus inverted microscope equipped with a Cascade II 2014 EM-CCD camera. Images were deconvolved with Applied Precision’s softWorx software. The images are shown as maximum intensity Z projections. An OlympusBX41 equipped with a DMK23U274 (ImagingSource®, Bremen, Germany) camera was also used, and image acquisition was through the software IC Capture 2.4. Super-resolution images were obtained with a Deltavision OMX microscope, a Structured Illumination (SIM) super-resolution microscope based on the Olympus GE Deltavision core equipped with a PCO Edge 5.5 sCMOS 2560x2160 (frame resolution, only 1024x1024 or lower) camera.

Confocal acquisitions in *Paramecium* cells were made with a Leica SP8 equipped with a UV diode (line 405) and three laser diodes (lines 488, 552, and 635) for excitation and two PMT detectors. For U-ExM data, images were collected with an inverted confocal Microscope Leica DMI 6000 CS. 3D stacks were acquired with 0.12 µm z-intervals and an x, y pixel size of 35 nm.

In all the cases, images were processed with ImageJ Fiji software (NIH, USA) and Photoshop CS5.

### 10 Electron microscopy

To visualize ultrastructure in *Paramecium,* the control and cells depleted of TBCCD1 using RNAi were fixed in 1% (v/v) glutaraldehyde and 1% (v/v) OsO_4_ in 0.05 M cacodylate buffer, pH 7.4 for 30 min. After washing the cells with cacodylate buffer, the cells were embedded in 2% agarose. Agarose blocks were then dehydrated in graded series of ethanol and propylene oxide and embedded in Epon. All ultrathin sections were contrasted with uranyl acetate and lead citrate. The sections were examined with a Jeol transmission electron microscope1400 (at 120 kV)

### 11 Image quantification and statistical analysis

Images were analyzed with the ImageJ Fiji software (NIH, USA). For the quantification of centriolar satellites, co-localization images were analyzed with the ImageJ Fiji software (NIH, USA) using ComDet (https://github.com/ekatrukha/ComDet) plugin. For the quantification of mean fluorescence intensity, CellProfiler v4.2.1 was used (McQuin et al. 2018). Statistical analysis was performed using Sigmastat v4.0.

**Legend - movie S1**- Mitotic RPE-1 cell constitutively expressing TBCCD1-GFP. Images were obtained at 2 min intervals for 8 h on DeltaVision Core System (Applied Precision) equipped with a climate chamber using a 60X objective (Olympus U-PLlan Apo N, UIS2, 1-U2B933). The movies were assembled using SoftWorx software. TBCCD1 was recruited from the cytoplasm to the midzone of the mitotic spindle of a dividing cell, where it binds to small segments of spindle MTs.

**Supplementary Figure S1 – Characterization of RPE-1 TBCCD1-GFP cell line. I.** Western blot for soluble and insoluble protein extracts of RPE-1 cell lines constitutively expressing TBCCD1-GFP (clone 6 and clone 5) or GFP alone (clone 10). Antibody against GFP was used. Bands observed present a predicted molecular mass consistent with the respective proteins (TBCCD1-GFP for clones 6 and 5 and GFP for clone 10) molar masses. TBCCD1-GFP expressed by clone 5 RPE-1 TBCCD1-GFP cell line responds to the transfection of TBCCD1 siRNAs as previously published for endogenous TBCCD1 (Gonçalves et al. 2010). The commercial antibody against TBCCD1 used in these studies recognizes both the endogenous and the TBCCD1-GFP forms in total protein extracts of RPE-1 cells. **II.** Immunofluorescence microscopy images for RPE-1 TBCCD1-GFP clone 5 cell line. **A** and **B** - TBCCD1-GFP (green), Centrin (red) was used as a centriolar marker, and DAPI shows DNA stained in blue. Arrows indicate TBCCD1-GFP at the centrosome and arrowheads at midzone MTs. **C** - TBCCD1-GFP (green), α-tubulin (red), and DAPI shows DNA stained in blue. Scale bars = 10 μm. **III.** Immunofluorescence microscopy images from frames of dividing clone 5 RPE-1 TBCCD1-GFP cell live imaging. Images show TBCCD1-GFP throughout cell division. TBCCD1-GFP is always present at the centrosomes and, during mitosis, is recruited from the cytoplasm to the midzone and later to the midbody, where it concentrates.

**Supplementary Figure S2 – Optimization of Taxol concentration for microtubule stabilization experiments.** Immunofluorescence images for α-tubulin (green) and DAPI showing DNA stained in blue. Cells were exposed to different concentrations of Taxol (0.1 nM and 0.5 nM) for 24 h to test the best concentration to avoid excessive alteration of cell and nucleus morphology and the formation of MTs bundles. Scale bars = 10 μm.

**Supplementary Figure S3. A-F-** Impact of TBCCD1 depletion in different centriolar DA and SDA proteins. Z projections of expansion microscopy combined with super-resolution 3D SIM images for the DA protein CEP164 (green) and SDA proteins ODF2 (red) and CEP128 (red) in RPE-1 cells (**A, C, and E**) and RPE-1 cells subjected to TBCCD1 specific siRNAs (**B, D, and F**). Centriole MTs were stained with an antibody against acetyl-α-tubulin (in green, except in A and D, that is in red). TBCCD1 depletion did not affect the localization of CEP164, ODF2, and CEP128. Scale bars = 0.5 μm.

**Supplementary Figure S4- I.** Schematic representation of the functional domains ascribed to human TBCC, RP2, TBCCD1, and *Paramecium* TBCCD1. **II.** The *Paramecium* protein presents the two characteristic functional domains of TBCCD1: the CARP domain and the TBCC domain. Sequence alignment of the putative TBCCD1 amino acid residues sequence of humans with the two TBCCD1 proteins of *Paramecium*. A comparison between human and Paramecium’s putative TBCCD1 amino acid sequence revealed 24.04 % identity. These values increase to 25.6% at the TBCC domain. The two *Paramecium* putative TBCCD1 amino acid residues sequence share 92.1% of identity. Highlighted in yellow are the amino acid residues of the TBCC domain. Highlighted in red wavy underline are the amino acid residues of the CARP domain. Highlighted in blue double underline are the siRNA target sequences in *Paramecium*. **III.** Decrease of the GFP signal in *Paramecium* TBCCD1-GFP transformants after TBCCD1 depletion compared to control cells. **A**- Differential interference contrast microscopy image (DIC) of a mixture of *Paramecium* cells corresponding to the same cells in image B. *Paramecium* cells treated with RNAi specific for TBCCD1 were fed with Indian ink allowing the visualization of feeding vacuoles (arrows) and to distinguish them from transformants that only express TBCCD1-GFP (controls). Both types of cells were mixed. **B**- Z projections of confocal sections at the ventral surface of a mixture of transformants expressing TBCCD1-GFP (control) and *Paramecium* transformants expressing TBCCD1-GFP treated with RNAi specific for TBCCD1 (show Indian ink in vacuoles in A). TBCCD1-GFP accumulates in vacuoles of control cells. In cells treated with RNAi specific for TBCCD1, the fluorescence disappeared, and BBs were only observed due to the staining with the antibody against poly-glutamylated tubulin (red, ID5). The results show that feeding the cells with TBCCD1-specific dsRNAs caused the complete depletion of TBCCD1-GFP. Scale bar = 20 μm.

**Supplementary Figure S5 – Optimization of flag-BirA*-TBCCD1 induced expression by tetracyclin in HEK293T Flp-In-Trex Cells.** Immunofluorescence images of HEK293T Flp-In-Trex cells and HEK293T Flp-In-Trex flag-BirA*-TBCCD1 cells. **A-D** – Different times of flag-BirA*-TBCCD1 expression induced by 0.5 μg/mL tetracycline in HEK293T Flp-In-Trex cells. γ-tubulin (green) is used as a centrosome marker, flag-BirA*-TBCCD1 (red), and DAPI shows DNA stained in blue. Arrowheads in merge show flag-BirA*-TBCCD1 staining centrosomes. Scale bars = 10 μm. **E-G** - Immunofluorescence images of biotinylated proteins in HEK293T Flp-In-Trex cells, HEK293T Flp-In-Trex flag-BirA* cells, and HEK293T Flp-In-Trex flag-BirA*-TBCCD1 cells induced by 0.5 μg/mL tetracycline and in the presence of 50 μM biotin. Cells were incubated with an antibody against the *flag* tag (green) to visualize the tagged expressed proteins and with Streptavidin conjugated with a fluorochrome (red) to allow visualization of biotinylated proteins. DAPI shows DNA stained in blue. In the merge, arrowheads show centrosomes with biotinylated proteins and tagged proteins. Scale bars = 10 μm.

**Supplementary Figure S6 – Validation of flag-BirA*-TBCCD1 induced expression and biotinylation of proteins by Western Blot analysis.** Total protein extracts were prepared from HEK293T Flp-In-Trex cells, HEK293T Flp-In-Trex flag-BirA* cells, and HEK293T Flp-In-Trex flag-BirA*-TBCCD1 cells and analysed by 10% SDS-PAGE followed by Western blot. **A**- Using the antibody anti-flag M2 is only possible to detect the expression of flag-BirA* in HEK293T Flp-In-Trex flag-BirA* cells induced by 1 μg/mL tetracycline; (+) – tetracyclin present; (-) - tetracyclin absent. **B** - Validation of protein biotinylation by flag-BirA* in HEK293T Flp-In-Trex flag-BirA* cells. Using HRP conjugated streptavidin, a different pattern of protein biotinylation is only detected when HEK293T Flp-In-Trex flag-BirA* cells were induced by 1 μg/mL tetracycline in the presence of 50 μM biotin; (+) – biotin present; (-) – biotin absent. **C** - Validation of protein biotinylation by flag-BirA*-TBCCD1 in HEK293T Flp-In-Trex flag-BirA* cells. Using HRP conjugated streptavidin, a specific pattern of protein biotinylation is only detected when HEK293T Flp-In-Trex flag-BirA*-TBCCD1 cells were induced by 1 μg/mL tetracyclin and 50 μM biotin was present. The induced expression of flag-BirA*TBCCD1 by tetracyclin in HEK293T Flp-In-Trex flag-BirA*-TBCCD1 cells was only observed using the antibody anti-flag M2 only when the cells were induced by tetracycline. The detected protein molecular mass is compatible with flag-BirA*-TBCCD1 protein size. **D** – Silver stained SDS-PAGE analysis for Streptavidin immunoprecipitated biotinylated proteins extracted from HEK293T Flp-In-Trex, HEK293T Flp-In-Trex flag-BirA* and HEK293T Flp-In-Trex flag-BirA*-TBCCD1 cells. The pattern of immunoprecipitated biotinylated proteins in HEK293T Flp-In-Trex flag-BirA*-TBCCD1 cells is different from that of total extracts (T), soluble fraction extracts (SF) or insoluble fraction (IF) obtained from the other cell lines used. Black arrows represent the visible differences between the soluble and insoluble fractions of the extracts.

**Supplementary Figure S7- A-** Protein-protein associations determined by STRING (https://string-db.org/) showing the relationships between centrosomal proteins selected from the TBCCD1 proximity network with a centriolar distal localization. Red line - experimentally determined; Green line - gene neighborhood; Blue line - co-expression. **B-** GO enrichment analysis for Cellular Components was performed using the Gene Ontology Resource (http://geneontology.org/). X-axis presents the fold enrichment for the Cellular Components GO term; Y-axis presents the -log P for the False Discovery Rate (FDR). Circle size is proportional to the number of proteins of the TBCCD1 interactome associated with each GO term. Only terms with a Fold enrichment > 5 and a -log P (FDR) > 5 are presented.

