## Supplementary figures and images for "The maintenance of centriole appendages and motile cilia basal body anchoring relies on TBCCD1"

### Supplemental Figure 1

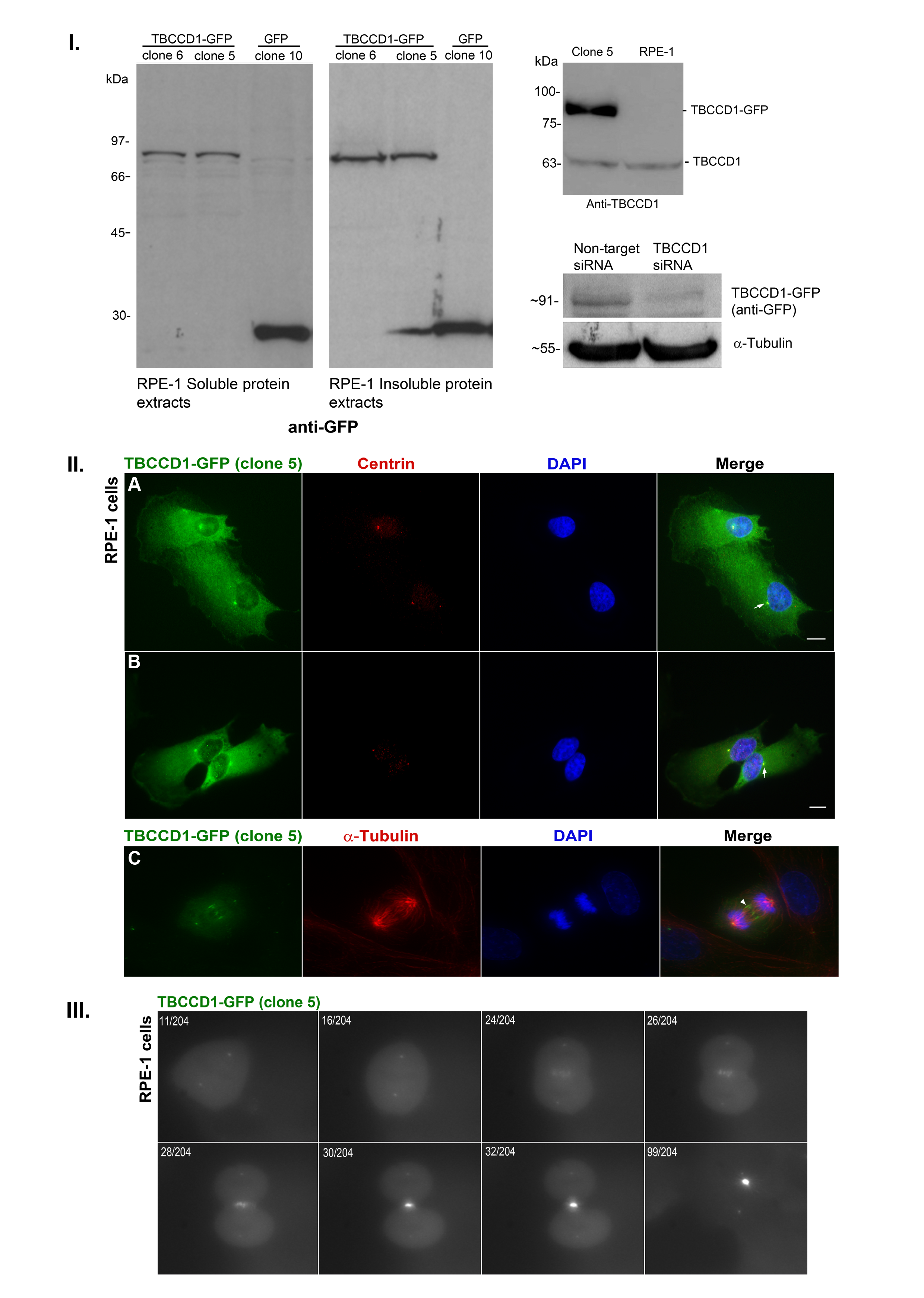

### Supplemental Figure 2

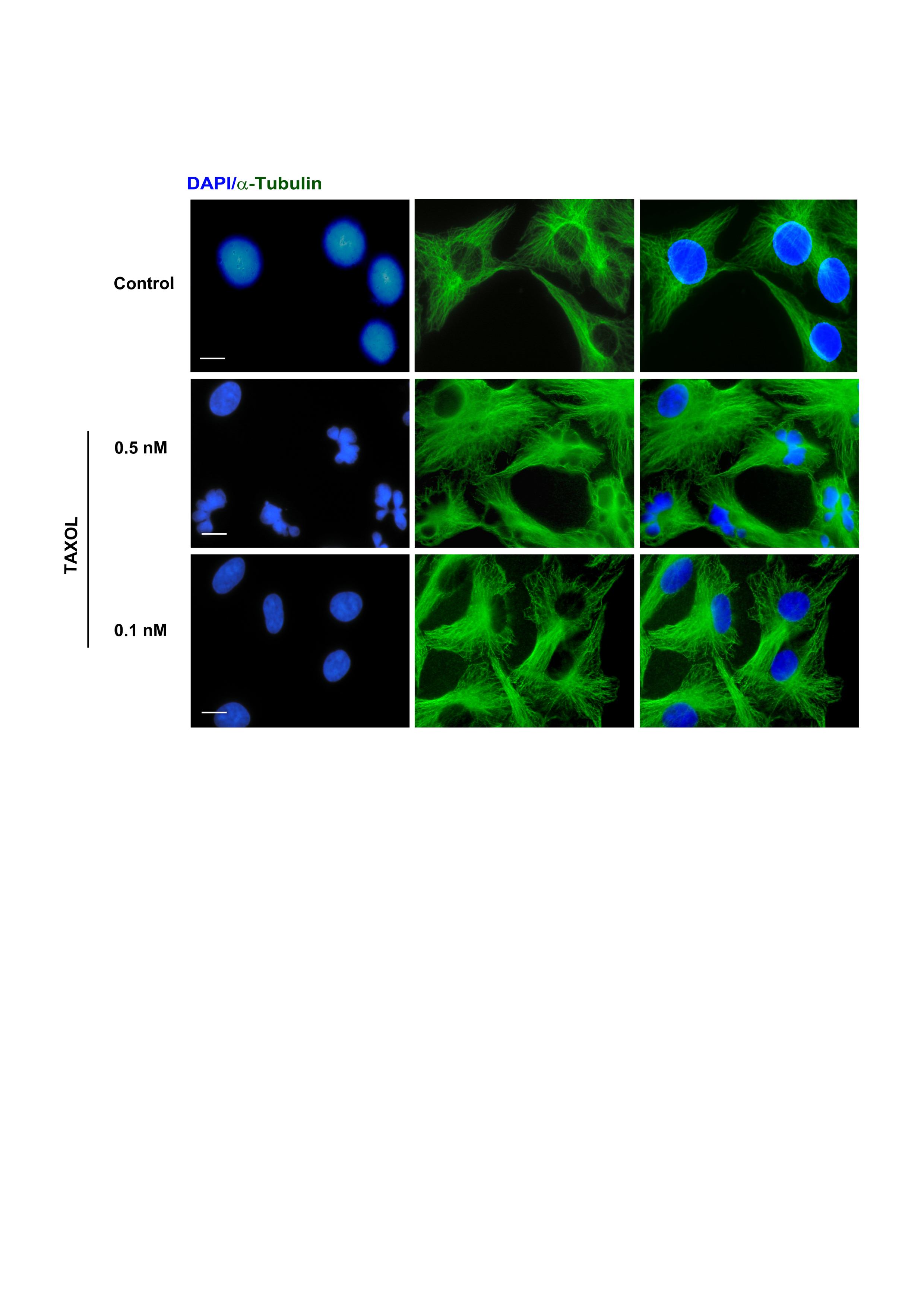

### Supplemental Figure 3

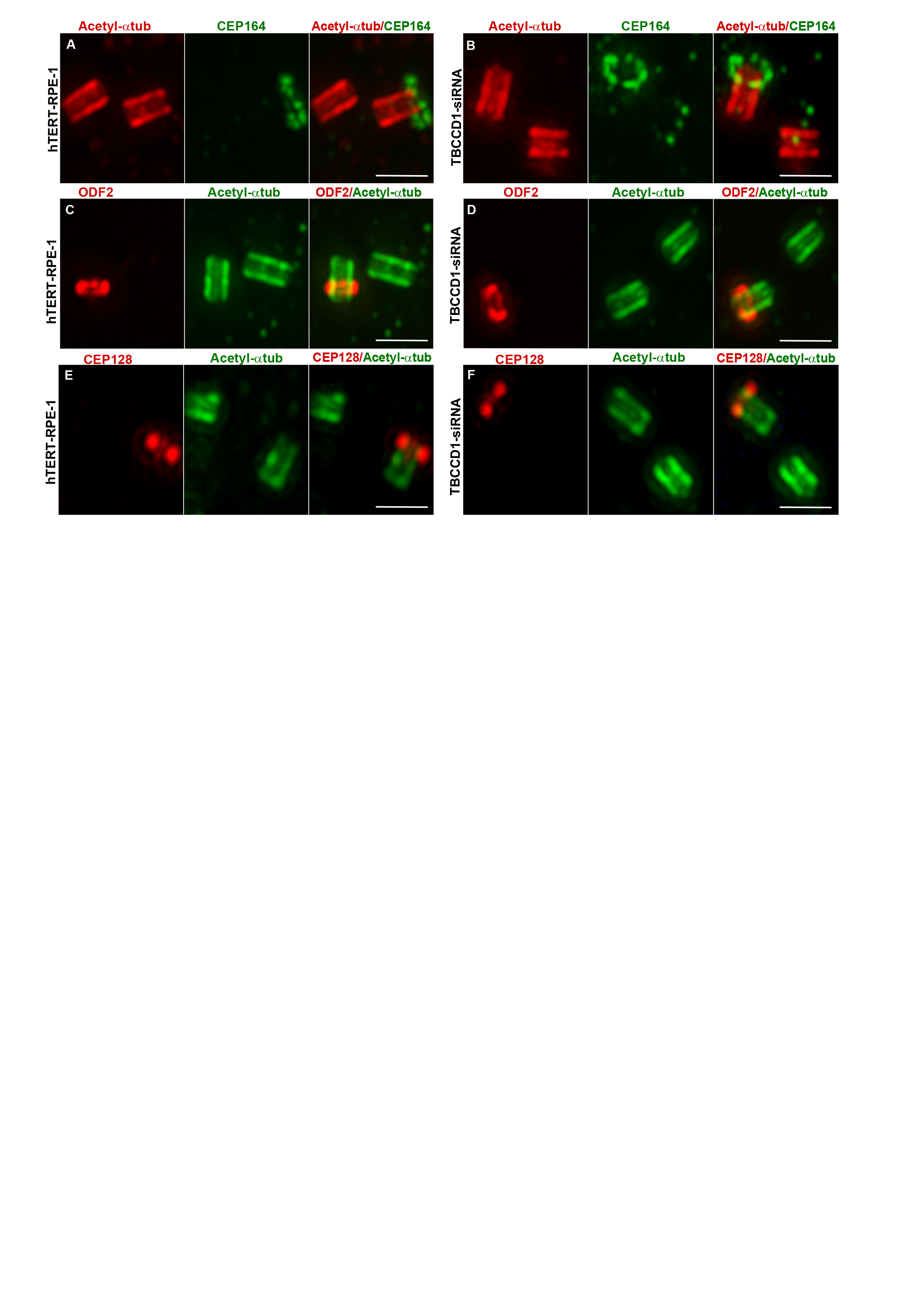

### Supplemental Figure 4

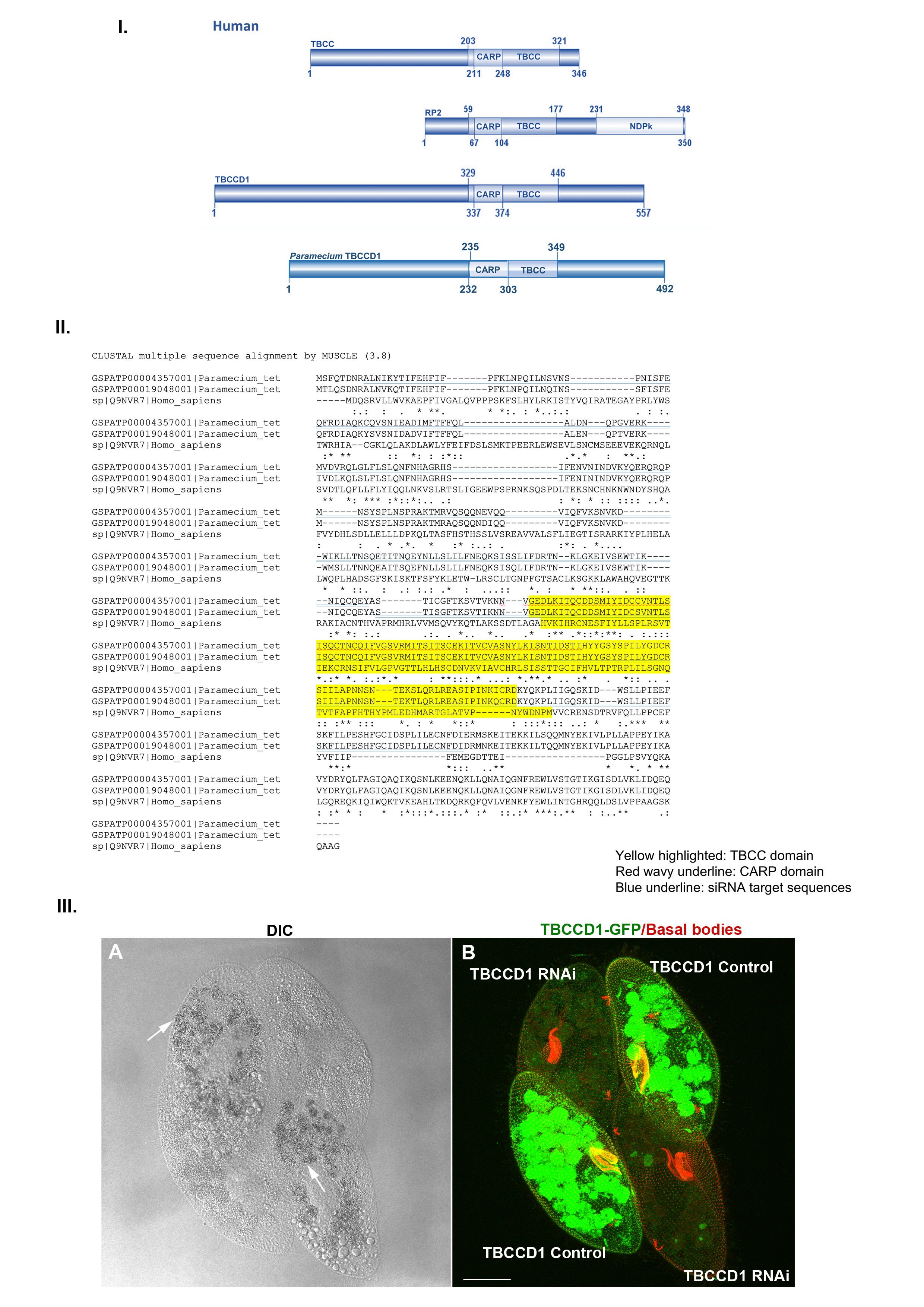

### Supplemental Figure 5

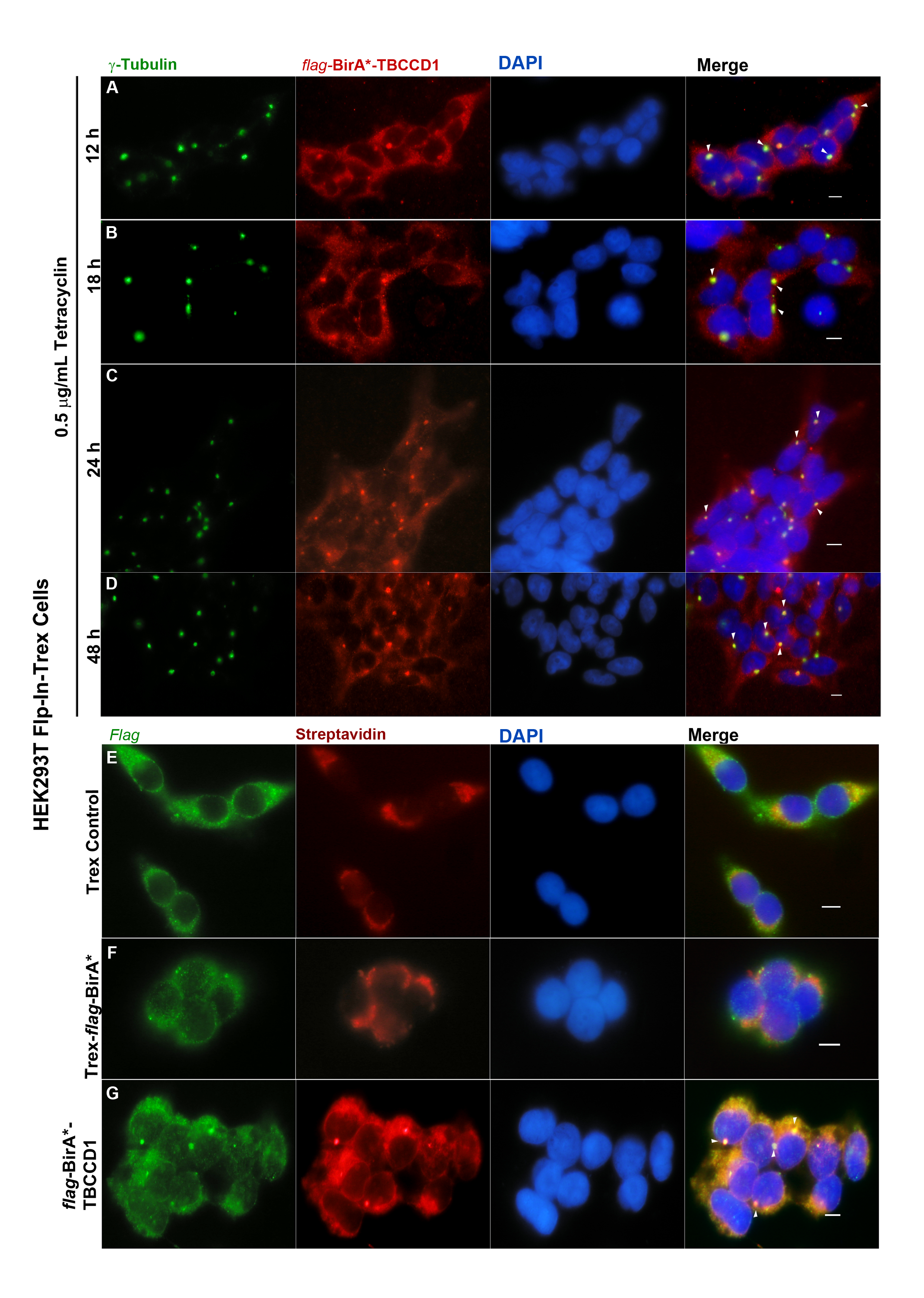

### Supplemental Figure 6

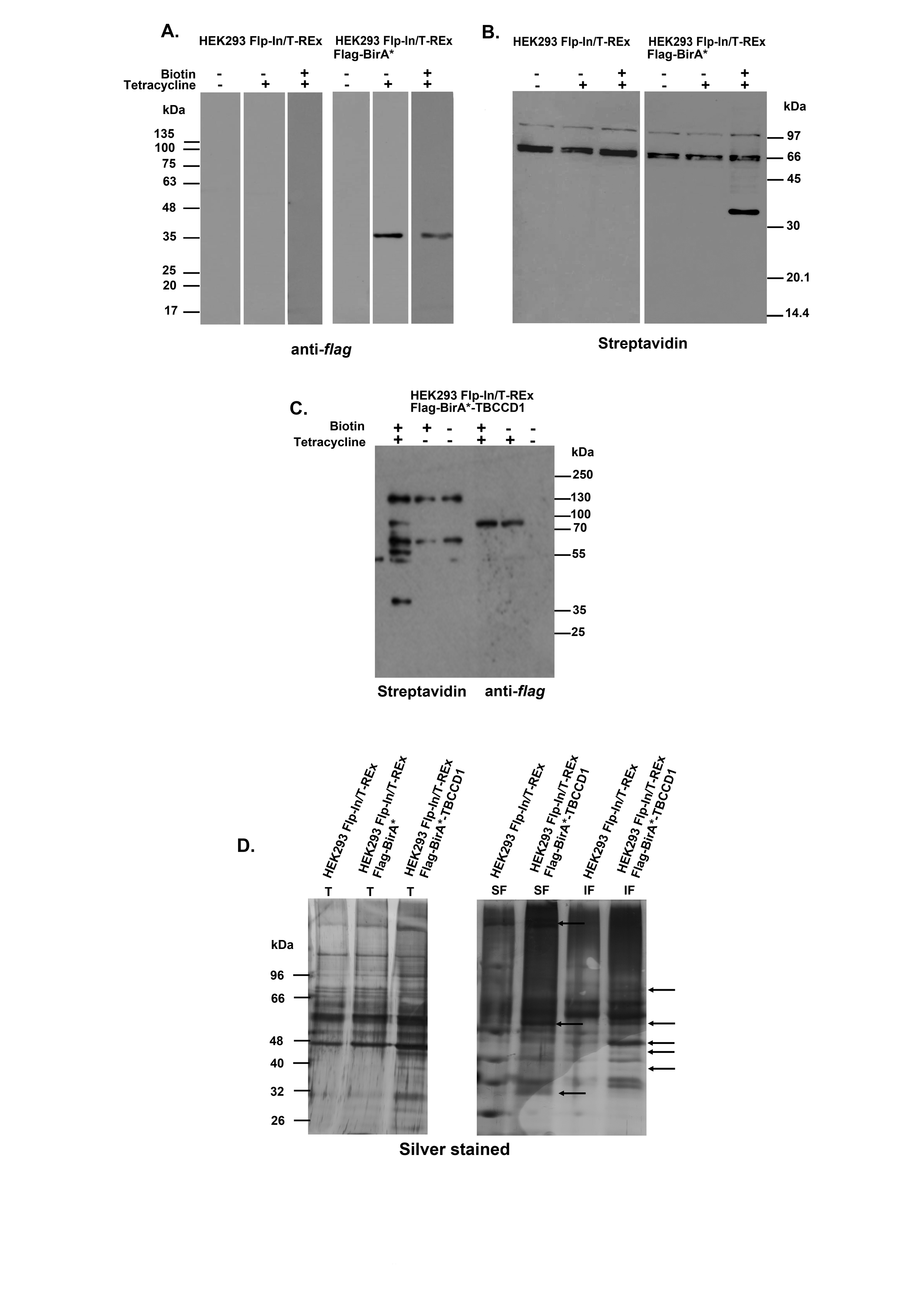

### Supplemental Figure 7

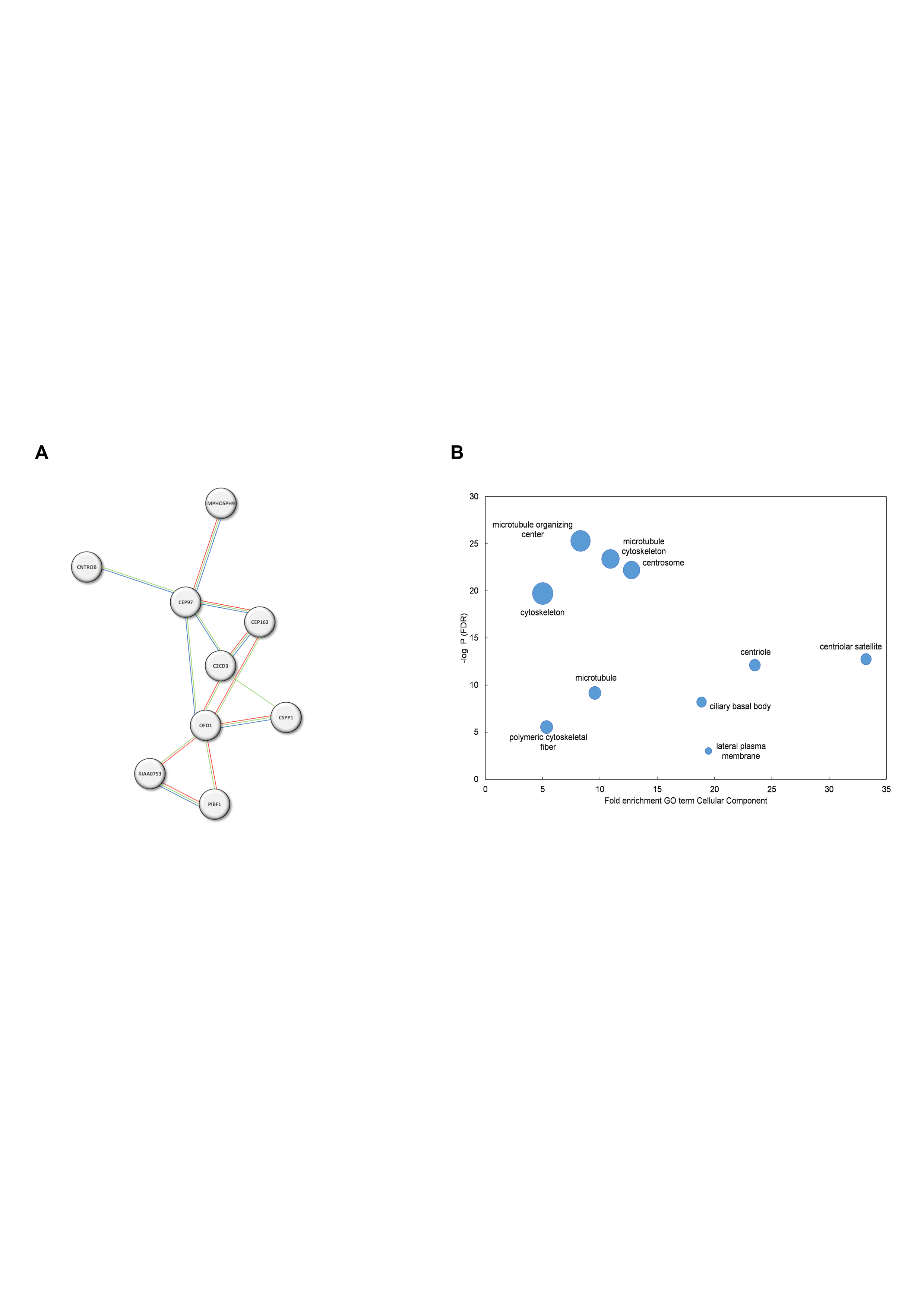
